# Senescent cell clearance ameliorates temporal lobe epilepsy and associated spatial memory deficits in mice

**DOI:** 10.1101/2024.07.30.605784

**Authors:** Tahiyana Khan, David J. McFall, Abbas I. Hussain, Logan A. Frayser, Timothy P. Casilli, Meaghan C. Steck, Irene Sanchez-Brualla, Noah M. Kuehn, Michelle Cho, Jacqueline A. Barnes, Brent T. Harris, Stefano Vicini, Patrick A. Forcelli

## Abstract

Current therapies for the epilepsies only treat the symptoms, but do not prevent epileptogenesis (the process in which epilepsy develops). Many cellular responses during epileptogenesis are also common hallmarks of *cellular senescence*, which halts proliferation of damaged cells. Clearing senescent cells (SCs) restores function in several age-associated and neurodegenerative disease models. It is unknown whether SC accumulation contributes to epileptogenesis and associated cognitive impairments. To address this question, we used a mouse model of temporal lobe epilepsy (TLE) and characterized the senescence phenotype throughout epileptogenesis. SCs accumulated 2 weeks after SE and were predominantly microglia. We ablated SCs and reduced (and in some cases prevented) the emergence of spontaneous seizures and normalized cognitive function in mice. Suggesting that this is a translationally-relevant target we also found SC accumulation in resected hippocampi from patients with TLE. These findings indicate that SC ablation after an epileptogenic insult is a potential anti-epileptogenic therapy.

## Introduction

Epilepsy is a chronic neurological condition characterized by recurrent seizures, affecting approximately 65 million individuals^1^. Temporal lobe epilepsy (TLE), characterized by seizures originating in the hippocampus and associated cortices, is one of the most prominent acquired epilepsies^2^. Current therapies for the treatment of epilepsy only control symptoms, but do not prevent epilepsy development (epileptogenesis) after injury or modify the disease state. Anti-epileptogenic therapies are thus a large unmet need. Cellular senescence, a conserved cellular program which halts cellular proliferation and upregulates pro-inflammatory factors seen in epilepsy in response to damage, may represent such a promising target.

Damage to the temporal lobe triggers an *epileptogenic cascade* which ultimately leads to spontaneous recurrent seizures (SRS),^3^ which often arise after a latent period of months to years in humans (days to weeks in rodents)^4^. Several processes are thought to contribute to epileptogenesis, including neuron loss, aberrant neuroplasticity, reactive gliosis, inflammation, and neurogenesis^5–9^. Acute changes following status epilepticus (SE) in experimental TLE models include hippocampal DNA damage^10,11^, excitotoxicity, and neuronal apoptosis^12–15^. Additionally, robust gliosis and increased production of proinflammatory cytokines and matrix metalloproteinases (MMPs) is evident following acute SE and persist throughout the latent and chronic periods^16–20^. In the chronic period, subsequent SRS may further exacerbate neurodegeneration, inflammation, and impair temporal lobe dependent cognitive functions^9^.

Chronic inflammation, gliosis, and excitotoxicity may contribute to accelerated brain aging in TLE, and this, in turn, may worsen cognitive function. Patients with TLE have older computed “brain ages” based on structural and functional MRI studies, with accelerated aging on the order of 4.5 to 11 years^21–23^. SC accumulation is frequently observed in normal aging, suggesting that cellular senescence is involved in age-associated cognitive impairments and neurodegenerative diseases^24–28^. With progressive age or precipitating injuries, cellular functions decline, senescence processes upregulate, and SC accumulation disrupts tissue structure and function^29^. Overall, the pathophysiological sequalae following epileptogenic insult and associated cognitive impairments point towards involvement of senescence pathways in epileptogenesis.

SCs play important physiological roles during wound healing, tumor suppression, and embryonic development^29^. SCs halt cell cycle progression by activating tumor suppressor pathways such as p53/p21 and p16^Ink4a^/Rb. SCs suppress apoptotic pathways but remain metabolically active and aberrantly influence the surrounding microenvironment by secreting pro-inflammatory cytokines, chemokines, MMPs, and growth factors, collectively termed as the senescence associated secretory phenotype (SASP)^30^. Many of these same molecules are upregulated after epileptogenic insults^16,17,20,31,32^.

In animal models of neurodegenerative disorders, selective SC clearance improved cognitive decline, normalized inflammation, and reduced neuropathology^26,27,33–37^. Given the advanced brain aging observed in TLE, the pro-inflammatory phenotype and reactive gliosis that are characteristic of the SASP component in cellular senescence (and have been proposed to contribute to the development of SRS), makes cellular senescence an intriguing target to prevent epileptogenesis. However, cellular senescence in epilepsy remains almost wholly unexplored^38^. Here we examined (1) the induction of cellular senescence and (2) the impact of SC clearance on epilepsy and cognitive comorbidities in experimental TLE. We found a robust upregulation of senescence-like cells in a mouse model of TLE (pilocarpine induced SE) and in resected hippocampi from individuals with TLE. SC clearance, using genetic and pharmacological ablation strategies, protected animals from epilepsy development, reduced SRS, and normalized cognitive phenotypes. These data indicate a causal role for SCs in the emergence of SRS after SE and suggest that SC clearance is a promising anti-epileptogenic therapeutic approach.

## Results

### Emergence of a microglial senescence phenotype during the latent phase of epileptogenesis

We characterized a senescence-like phenotype throughout epileptogenesis in the well-established pilocarpine-induced mouse model of experimental TLE. Systemic pilocarpine induces SE (an acute epileptogenic insult), followed by a seizure free latent period after SE termination (2-4 weeks), pathological reorganization in the brain, and the emergence of spontaneous recurrent seizures and cognitive impairment^39,40^. Mice were monitored for 2 hours after pilocarpine administration (280mg/kg), after which seizures were terminated with diazepam (5mg/kg). Onset of SE was defined as 5 min of uninterrupted bilateral forelimb clonus (Stage 3 seizures). We employed orthogonal strategies to assess the extent of a senescence-like phenotype with widely utilized putative senescence markers: p16 (*cdkn2a*), p21 (*cdkn1a*), and senescence-associated beta-galactosidase (SA-βgal) and compared their expression in mice that underwent SE to seizure-naïve, age matched controls (No SE). p16 and p21 are tumor suppressor proteins that inhibit cell cycle progression and are associated with cellular senescence, while β-gal is an enzyme used as a biomarker to detect senescent cells by increased lysosomal activity^29^. p16 and p21 mRNA expression was significantly increased 2 weeks following SE in the hippocampus. This time point falls well within the typical latent period in this model. p16 and p21 mRNA expression was also significantly increased in the chronic phase (8 weeks after SE) (Fig. 1A-E). Additionally, we histologically examined p16 expression in a p16-reporter mouse line. In this model, tandem-dimer Tomato (tdTom) is knocked into exon 1α of the p16^INK4a^ locus which results in tdTom expression following activation of the p16 promoter^41^. Increased p16-tdTom expression was also observed 2 and 8 weeks after SE (Fig. 1F-G). We found a similar pattern of p16 expression with p16 antibody staining, validating the use of the p16 antibody immunolabeling (Supplementary Fig. 1). Moreover, these experiments revealed a delayed upregulation of p16, after 2 weeks that was not apparent on day 1 following SE. Unlike p21 mRNA expression, p21 protein expression was significantly increased only at the 8 week post SE timepoint (Fig. 1H-I). Lastly, we found a significant increase in SA-βgal expression at 2 and 8 weeks after SE (Fig. 1J-K). Our results indicate an increase in a senescent-like phenotype following SE that arises during the latent period and continues to increase after chronic epilepsy is established.

**Fig. 1.**
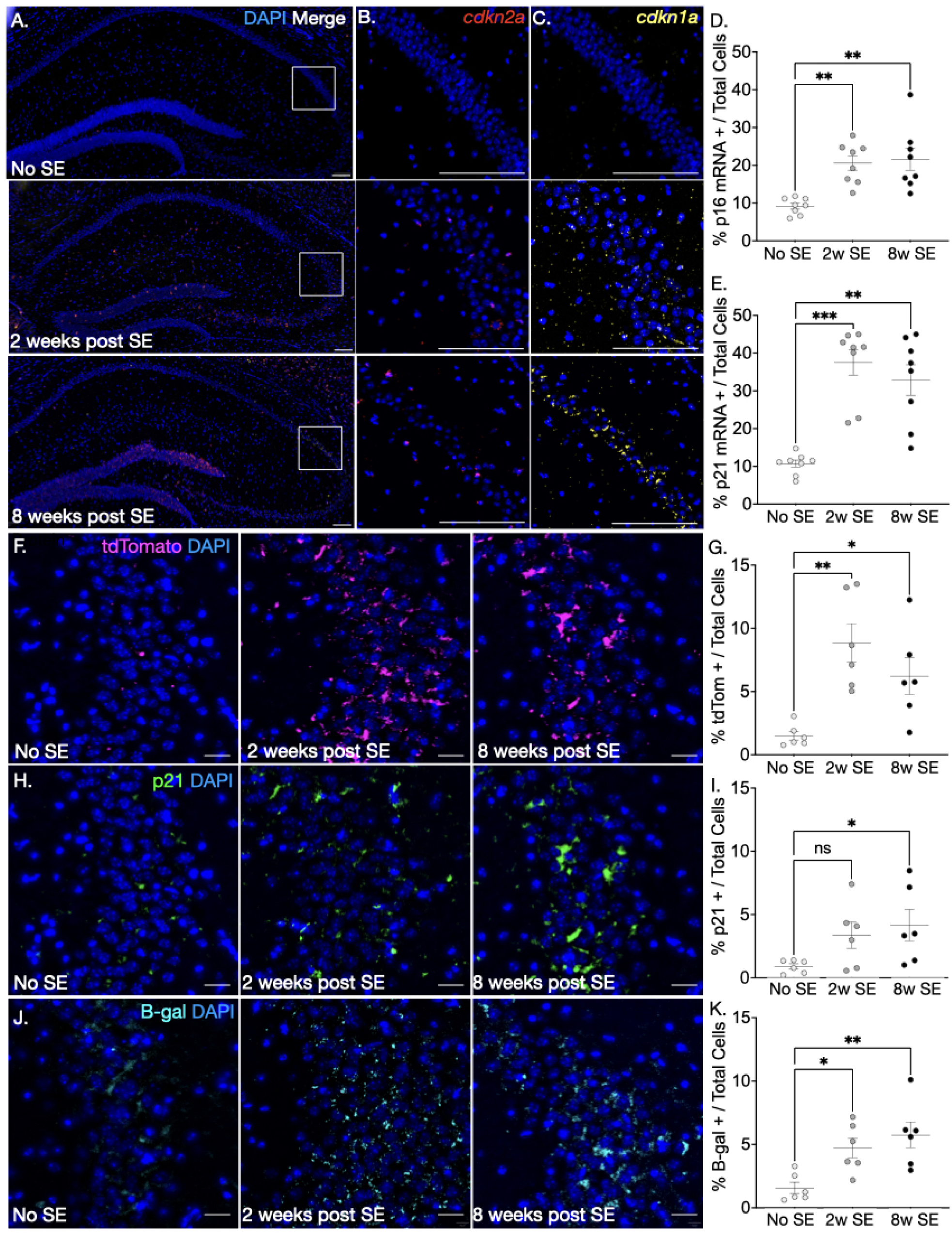
Characterization of the senescence phenotype following an epileptogenic insult. (A.) Representative hippocampal images of *cdkn2a* and *cdkn1a* mRNA transcripts of No SE controls (top), 2 weeks post SE (middle), and 8 weeks post SE (bottom). All zoomed in images represent a similar region of interest. Zoomed in representative images for (B.) *cdkn2a* mRNA (red) (C.) *cdkn1a* mRNA (yellow). (D.) Percent of *cdkn2a* mRNA positive cells over total cells in the hippocampus (p=0.0005, H=15.37. No SE vs. 2w SE: p=0.0012, Z=3.429; No SE vs. 8w SE: p=0.0016, Z=3.359) (E.) Percent of *cdkn1a* mRNA positive cells over total cells in the hippocampus (p=0.0004, H=15.77; No SE vs. 2w SE: p=0.0004, Z=3.712; No SE vs. 8w SE p=0.0042, Z=3.076). Representative zoomed CA3 images from No SE (left), 2 weeks post SE (middle), 8 weeks post SE (right) conditions for (F.) p16 immunofluorescence with tdTomato reporter (magenta, top), (H.) p21 immunofluorescence (green, middle), and (J.) SA-βgal (cyan, bottom). (G.) Quantification of percent tdtTomato positive cells from total detected cells in the hippocampus (p=0.0010, H=10.89. No SE vs. 2w SE: p=0.0028, Z=3.190; No SE vs. 8w SE: p=0.0401, Z=2.325). (I.) Quantification of percent p21 positive cells from total detected cells in the hippocampus (p=0.0568, H=5.535. No SE vs. 2w SE: p=0.1668, Z=1.731; No SE vs. 8w SE: p=0.0495, Z=2.245). (K.) Quantification of percent SA-βgal positive cells from total detected cells in the hippocampus (p=0.0055, F(2,15)=7.520. No SE vs. 2w SE: p=0.0221, Z=2.541; No SE vs. 8w SE: p=0.0099, Z=2.812). D-E: n=8 hippocampi were analyzed per group. G, I, K: n=6 animals per group. For all: Kruskal-Wallis Test, Dunn’s multiple comparison test. Mean = +/- S.E.M.

We next determined cell-type specificity of the senescent-like phenotype during chronic epilepsy (8+ weeks after SE). Using the p16-tdTom reporter strain, we assessed tdTom colocalization with markers for microglia (Iba1), astrocytes (GFAP), oligodendrocytes (Olig2), and immature neurons (DCX), and found tdTom colocalization predominantly with microglia (Fig. 2A, Supplementary Fig. 2). We found similar results when we replicated these findings with p16 antibody labeling (Supplementary Fig. 3). Additionally, p16 and p21 mRNA significantly colocalized with a microglial transcript (*aif1*) both at 2- and 8-weeks post SE (Fig. 2B-C, E-M). These results indicate microglia as the cell type that most robustly displays increases in senescence markers following SE. This is particularly interesting as microglia are increasingly recognized as a contributor to epileptogenesis^42–44^.

**Fig. 2.**
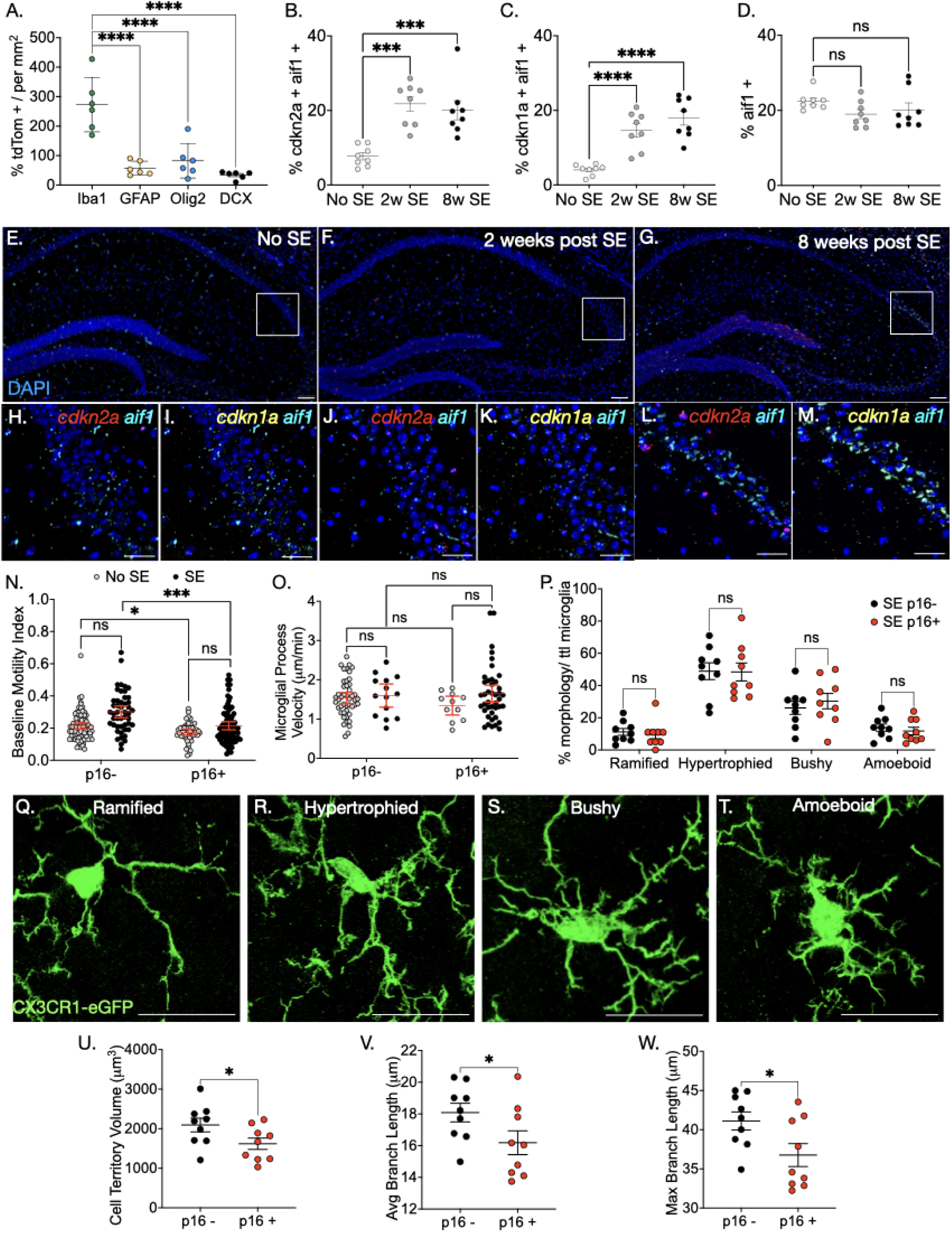
A senescence-like phenotype is predominantly expressed in microglia. (A.) Quantification of p16 tdTom reporter colocalization with microglia (Iba1), astrocytes (GFAP), oligodendrocytes (Olig2), and immature neurons (DCX) in the hippocampus 8 weeks post SE (n=6 per group). p<0.0001, F(2,21)=22.64. Iba1 vs. GFAP: p<0.0001, t=6.637, DF=20; Iba1 vs. Olig2: p<0.0001 t=5.860, DF=20; Iba1 vs. DCX: p<0.0001, t=7.354, DF=20). (B.) Quantification of percent of *cdkn2a* and *aif1* mRNA positive cells over total detected cells in the hippocampus. (p<0.0001, F_(2,21)_=15.19; No SE vs. 2w SE: p=0.0001, t=5.052, DF=21; No SE vs. 8w SE: p=0.0002, t=4.43, DF=21). (C.) Quantification of percent of *cdkn1a* and *aif1* mRNA positive cells over total detected cells in the hippocampus. (p<0.0001, F_(2,21)_=22.75; No SE vs. 2w SE: p<0.0001, t=4.934, DF=21; No SE vs. 8w SE: p<0.0001, t=6.450, DF=21). (D.) Quantification of *aif1* mRNA positive cells over total detected cells in the hippocampus. (p=0.2114, F_(2,21)_=1.675; No SE vs. 2w SE: p=0.1662, t=1.796, DF=21; No SE vs. 8w SE: p=2.428, t=1.202, DF=21). Representative hippocampal images of *cdkn2a* (red) and *cdkn1a* (yellow) mRNA multiplexed with *aif1* (cyan) mRNA of (E.) No SE, (F.) 2 weeks post SE, and (G.) 8 weeks post SE. Zoomed in representative image of *cdkn2a* and *aif1* probes (H.) No SE, (J.) 2w SE, (L.) 8w SE. Zoomed in representative image of *cdkn1a* and *aif1* probes (I.) No SE, (K.) 2w SE, (M.) 8w SE. (N.) Baseline Motility Index. Main effect of p16 expression (p<0.0001, F(1,213.759)=17.454), Interaction between p16 expression, disease conditions, and hippocampal subregion (p=0.02251, F(4, 213.841)=2.911). No main effect of disease condition (p=0.152, F(1,8.323)=2.484), or hippocampal subregion (p=0.193, F=2,214.538)=1.66). No SE-p16-vs. p16+: p=0.0262, t=2.29, DF=211.686; SE-p16-vs. p16+: p=0.00024, t=3.75, DF=215.743. p16-microglia: No SE (n=81), SE (n=48); p16+ microglia: No SE (n=45), SE (n=78). (O.) Microglial Process Velocity in response to 3mM ATP. No main effect of p16 (p=0.323), condition (p=0.460), or interaction of p16 x condition (p=0.689). p16-microglia: No SE (n=46), SE (n=14); p16+ microglia: No SE (n=11), SE (n=42). (P.) Percent of p16 positive and negative microglia per morphology category in SE hippocampi. Main effect of morphology (p<0.0001, F(3,72)=34.32; no main effect of p16 (p=0.9788, F(1,72)=0.0007131; no interaction effect of morphology and p16 (p=0.9520, F(3,72)=0.1133). Two-Way ANOVA, Holms-Šídák’s multiple comparison test. Representative CX3CR1-eGFP positive microglia for each morphology category from confocal z-stack images (Q.) Ramified, (R.) Hypertrophied, (S.) Bushy, (T.) Amoeboid. (U.) Cell territory volume of p16- and p16 + microglia in SE animals (p=0.0260, t=2.727, DF=8). (V.) Average branch length of p16- and p16 + microglia in SE animals (p=0.0374, t=2.491, DF=8). (W.) Average branch length of p16- and p16 + microglia in SE animals (p=0.0253, t=2.744, DF=8). B-D: n=8 hippocampi were analyzed per group. P,U-W: n=9 animals. A-D: Ordinary One-way ANOVA, Holms-Šídák’s multiple comparison test. N-O: Mixed Model Analysis; Šídák’s multiple comparison test. U-W: Paired t-test. Mean = +/- S.E.M.

Microglia constantly scan the extracellular environment through their processes (basal motility) and direct their processes towards potentially dangerous signals (through the activation of purinergic receptors), however, these processes are altered in pathological conditions^45–47^. Their “resting” state morphology is ramified with a small soma and fine cellular processes. In response to loss of CNS homeostasis, microglia take on an “activated” morphology in which cellular processes retract and the soma is enlarged to an amoeboid appearance^48^. There are several intermediate stages between the two extremes of “ramified/resting state” and “amoeboid/activated” microglia, such as “hypertrophied” and “bushy”^49^. We suspected that senescence processes may impact normal microglial function in epilepsy and thus investigated the properties of p16+ and p16-microglia. We crossed the p16-tdTom reporter with a microglia reporter line, CX3CR1^eGFP+^ ^50^, and examined baseline and responsive motility in acute hippocampal slices. As expected, p16+ microglia had reduced baseline spontaneous motility in both control (No SE) and chronic epilepsy (SE) conditions (Fig. 2N). We elicited responsive motility in acute slices by releasing 3mM ATP from a patch pipette and found no differences in microglial process velocity between chronic epilepsy mice and controls, nor between p16+ and p16-microglia (Fig. 2O; Supplementary Fig. 9). We did not observe changes in morphology between p16+ and p16-microglia in the hippocampus after epilepsy development (Fig. 2P-T). However, in *ex vivo* acute hippocampal slices of epilepsy and control conditions, we found a higher proportion of p16+ amoeboid microglia and reduced “resting state” p16-microglia in epilepsy (Supplementary Fig. 4). We further examined whether p16 expression influenced microglial properties after epilepsy development using a modified version of 3DMorph automated analysis^51^. p16+ microglia had reduced cell territory volume (Fig. 2U), reduced average (Fig. 2V) and maximum (Fig. 2W) branch lengths compared to p16-microglia, indicating a shift in morphology towards an activated state.

### p16-specific SC ablation throughout epileptogenesis rescues spatial memory impairments and reduces seizure burden

Due to the robust expression of a senescence-like phenotype, we hypothesized that clearance of SCs after an epileptogenic insult (SE) reduce seizure burden and cognitive comorbidities. We targeted p16-expressing cells with INK-ATTAC mice, which allows for inducible depletion of p16+ cells upon activation of a caspase transgene by a synthetic dimerizing agent, AP20187 (AP)^33^. Mice were subjected to SE, and randomly assigned to receive twice weekly 2mg/kg (i.p.) AP to eliminate p16+ cells (SE+ AP) or vehicle (SE + Veh) immediately following SE termination until the end of the experiment (Fig. 3A). We chose this treatment paradigm in our study to examine an *anti-epileptogenic* effect.

**Fig. 3.**
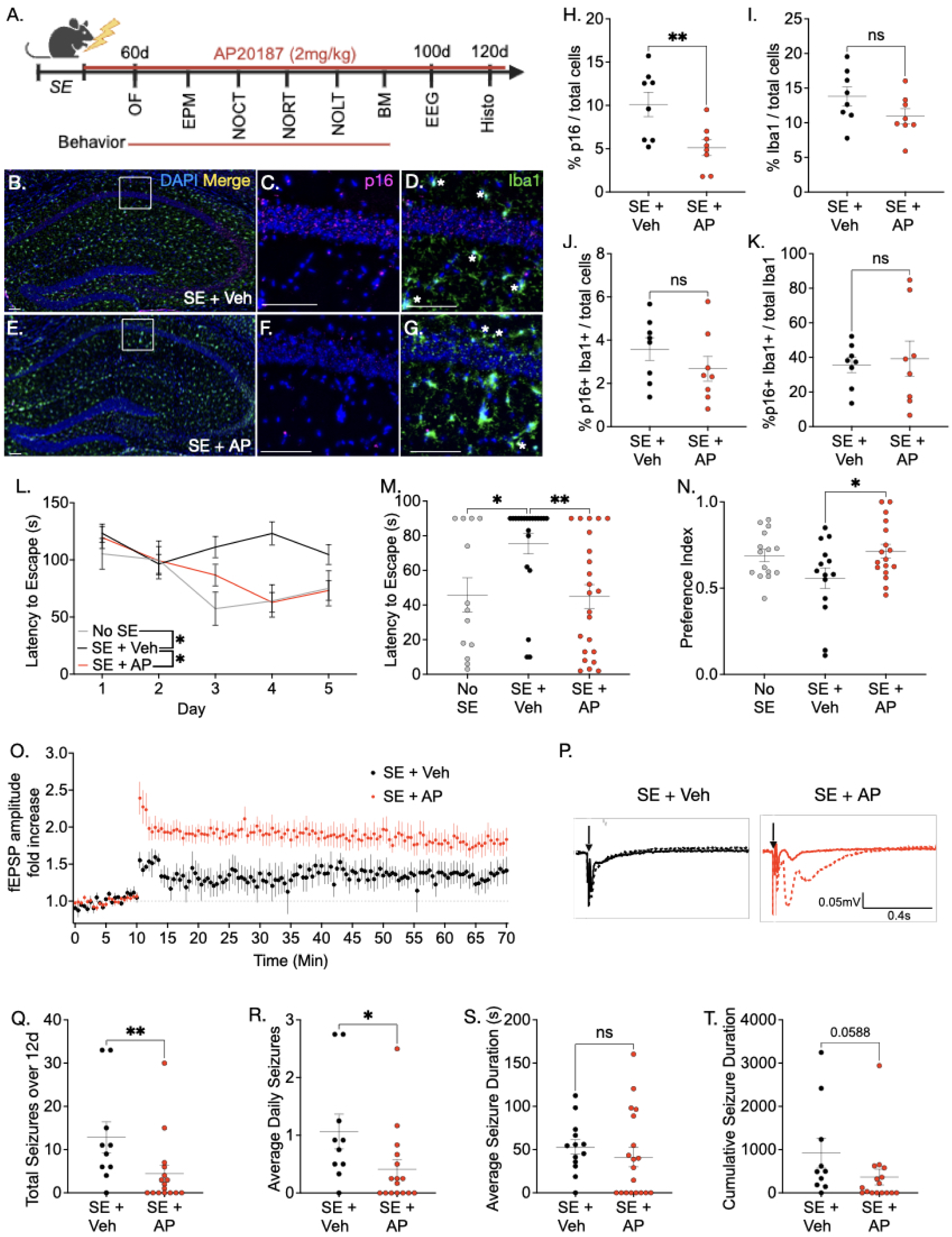
p16-specific senescent cell ablation attenuates seizure burden and spatial memory deficits following epilepsy development in INK-ATTAC. (A.) Experimental timeline. 4-month-old male and female mice receive vehicle or AP treatment following SE termination until the end of the experiment. 8 weeks after SE animals undergo behavioral tests, followed by 12 days of EEG implantation and EEG monitoring. At the end of the study, brains are processed for histological analysis of senescence markers. Representative image of p16 antibody (magenta) and Iba1 (green) expression in vehicle-treated (B.) and AP-treated mice (E.) 4 months following SE. Zoomed in representative image of p16 and merged immunofluorescence in vehicle-treated (C, D.) and AP-treated mice (F, G.). (H.) Percent p16 positive cells from total detected nuclei (p=0.0098, t=2.987, DF=14). (I.) Percent Iba1 positive microglia from total detected nuclei (p=0.119, t=1.656, DF=14). (J.) Percent p16 positive microglia from total detected nuclei in hippocampus (p=0.269, t=1.167, DF=14). (K.) Percent p16 positive microglia from total detected microglia (p=0.756, t=0.3172, DF=14). (L.) Barnes maze training (p<0.0001, F(2,265.66)=11.230 Mixed Model Analysis, Šídák’s multiple comparison test). Main effect of treatment (p<0.0001; F(2,265.66)=11.23), trial day (p=0.00054; F(4,96.62)=5.44), interaction between treatment and trial day (p=0.0398; F(8,96.62)=2.132); No SE vs. SE + Veh: p<0.0001; No SE vs. SE + AP: p=0.608; SE + Veh vs. SE + AP: p=0.0007. (M.) Barnes Maze Probe (p=0.0048, H=10.69, Kruskal-Wallis test); SE + Veh vs. No SE: p=0.0343, z=2.383; SE + Veh vs. SE + AP: p=0.0043, z=3.068; Dunn’s Multiple Comparison Test. (N.) Novel Location preference index after a 5 minute delay (p=0.046, F(2,43)=3.32; SE + Veh vs. No SE (p=0.053, t=1.99, DF=43), SE + Veh vs. SE + AP (p=0.037, t=2.45, DF=43; One-Way ANOVA, Holms-Šídák’s multiple comparison test). (O.) Long term potentiation (LTP) fEPSP amplitude fold increase in SE + Veh (black), and SE + AP (red). Main effect of condition (p=0.025, F(1,13)=6.39), time elapsed (p<0.0001, F(4.32,56.18)=9.38), and interaction between condition and time elapsed (p<0.0001, F(140,1820)=2.23). Two-Way ANOVA, Šídák’s multiple comparison test. (P.) Representative traces of SE + Veh (black) and SE + AP (red). (Q.) Total number of seizures over 12 days (p=0.0080, U=31.50). (R.) Average number of seizures per day (p=0.0136, U=34.50). (S.) Average duration of each seizure (p=0.153, U=86). (T.) Cumulative seizure duration (p=0.059, U=44.50). H-K: n=8 animals per group; L-M: no SE (n=13), Se + Veh (n=21), SE + AP (n=23). N: no SE (n=15), Se + Veh (n=14), SE + AP (n=17). N: Se + Veh (n=21), SE + AP (n=23); O-P: Se + Veh (n=4), SE + AP (n=11); Q-T: + Veh (n=10), SE + AP (n=16). H-K: Unpaired t-test; Q-T: Mann Whitney Test. Mean = +/- S.E.M.

We confirmed p16+ cell ablation in INK-ATTAC mice 4 months after SE (Fig. 3B-K). p16 expression was reduced by approximately 50% in the hippocampus in AP-treated mice (Fig. 3H). Studies have reported that a 30% reduction in putative senescence markers is sufficient for improvement in function^34,36,52–54^. Hippocampal microglial cell number was indistinguishable between vehicle and AP-treated groups (Fig. 3I) because p16-expressing microglia represent only a small fraction of the total hippocampal cell population (Fig. 3J) and in the microglial population (Fig. 3K). However, p16-specific cell elimination decreased protein expression of pro-inflammatory SASPs: IL-1β, CCL3, and CCL4, with a trend towards a reduction in TNF-α and CXCL9 (Supplementary Fig. 5). These findings suggest that p16-specific cell elimination in SE reduced the expression of p16+ cells in the hippocampus and SASP components without altering the overall microglial number.

We performed a battery of behavioral tests to assess whether ablating p16+ cells rescue cognitive functions following epilepsy development. Clearance of p16+ cells did not impact anxiety or exploratory behaviors (Supplementary Fig. 6A-D), or recognition memory functions (Supplementary Fig. 6F-I). However, spatial memory processes were rescued in the Barnes Maze (BM, Fig. 3L-M) and Novel Object Location test (NOLT; Fig. 3N). AP-treated SE mice and No SE mice learned to find the escape hole in the BM over time, whereas vehicle-treated mice did not learn throughout the training days (Fig. 3L). 24 hours after the last training day, mice were subjected to the probe test in which latency to find the escape hole was measured (Fig. 3M). AP-treated SE mice had a reduced latency to escape compared to vehicle-treated mice, indicating a rescue of spatial memory. 66.7% (14/21) of vehicle-treated mice failed to escape on the probe trial compared to 21.7% (5/23) of AP-treated mice (χ^2^ (1) = 9.03, p=0.0027), and 30.8% (4/13) of control mice (χ^2^ (1) = 4.15, p=0.041). We also examined spatial memory in NOLT, by measuring animals’ exploration of a familiar object moved to a new location compared to an unchanged object. AP-treated SE mice showed a higher preference for the novel location following a 5-minute delay period than vehicle-treated SE mice (Fig. 3N). To determine if p16-specific cell clearance altered hippocampal synaptic plasticity, we next examined Schaffer collateral to CA1 long term potentiation. In line with our behavioral findings, AP-mediated SC clearance resulted in an increase in long term potentiation (LTP) compared to vehicle-treated SE mice (Fig. 3O-P).

Lastly, we examined the impact of p16-positive cell ablation on seizure burden. SC clearance completely protected 40% of mice in the AP-treated group from spontaneous seizures. Moreover, AP-treated animals displayed significant reductions in the total number of seizures during the recording period (Fig. 3Q) and average number of daily seizures (Fig. 3R). Average seizure duration (Fig. 3S) and cumulative seizure duration (Fig. 3T) were not changed with SC elimination. These results indicate that p16-specific SC ablation throughout epileptogenesis rescues spatial memory deficits and seizure burden in chronic epilepsy.

### Global SC ablation with senolytics, D&Q, throughout epileptogenesis rescues spatial memory impairments and reduces seizure burden

While transgenic models employing p16-specific SC elimination provide mechanistic insight on senescence processes in various diseases, translatable strategies to clear SCs in patients are necessary for a therapeutic application. One approach takes advantage of SCs’ resistance to apoptosis and kills SCs through the Senescent Cell Anti-Apoptotic Pathway (SCAP) networks^55^. These include tyrosine kinase inhibitor, dasatinib, which promotes apoptosis by dependence receptors such as ephrins (in part by inhibiting Src kinase), and quercetin, a naturally occurring plant flavonoid^56^. Targeting SCs using senotherapies has been examined in several neurodegenerative disease models^25–27,35,37,57^. We were interested in assessing how targeting pro-survival pathways of SCs would impact epileptogenesis and cognitive functions, and how this SC ablation strategy may differ from p16-specific SC ablation. We utilized senolytic compounds, dasatinib and quercetin (DQ), following SE and throughout epileptogenesis to assess the resulting impact on seizure burden and cognitive impairments associated with TLE (Fig. 4A).

**Fig. 4.**
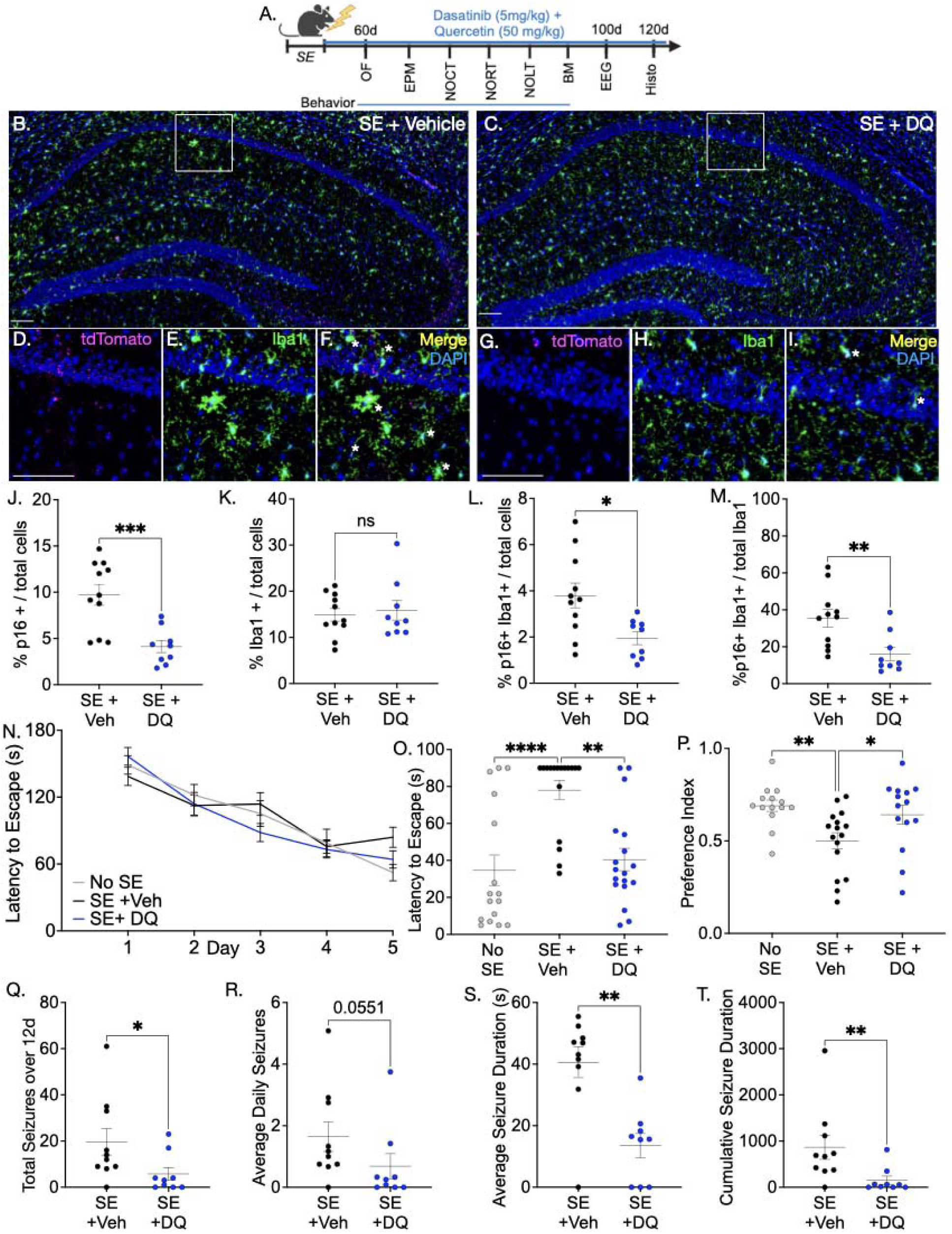
Senotherapy with DQ reduces seizure burden and spatial memory deficits following epilepsy development in C57BL/6 mice. (A.) Experimental timeline. 4-month-old male and female mice receive vehicle or DQ treatment following SE termination until the end of the experiment. 8 weeks after SE animals undergo behavioral tests, followed by 12 days of EEG implantation and EEG monitoring. At the end of the study, brains are processed for histological analysis of senescence markers. Representative image of p16 antibody (magenta) and Iba1 (green) expression in vehicle-treated (B.) and DQ-treated mice (C.) 4 months following SE. Zoomed in representative image of p16, Iba1, and merged immunofluorescence in vehicle-treated (D, E, F.) and DQ-treated mice (G, H, I.). (J.) percent p16 positive cells from total detected nuclei (p=0.0007, t=4.069, DF=18). (J.) percent Iba1 positive microglia from total detected nuclei (p=0.7004, t=0.3910, DF=18). (L.) percent p16 positive microglia from total detected nuclei in hippocampus (p p=0.0106, t=2.853, DF=18). (M.) percent p16 positive microglia from total detected microglia (p=0.0063, t=3.092, DF=18). (N.) Barnes maze training (p=0.592, F(2,233.129)=0.526; Mixed Model Analysis, Šídák’s multiple comparison test). Main effect of trial day (p<0.0001; F(4,96.932)=46.003, no main effect of treatment (p=0.592, F(2,233.129)=0.526), interaction between treatment and trial day (p=0.112, F(8,96.932)=1.682). (O.) Barnes Maze Probe (p<0.0001, H=19.44, Kruskal-Wallis test). SE + Veh vs. No SE: p<0.0001, z=4.17; SE + Veh vs. SE + DQ: p=0.0018, z=3.323; Dunn’s Multiple Comparison Test. (P.) Novel Location preference index after a 5 minute delay (p=0.0079, F(2,41)=5.454; SE + Veh vs. No SE (p=0.006, t=3.153, DF=41), SE + Veh vs. SE + DQ (p=0.0241, t=2.342, DF=41; One-Way ANOVA, Holms-Šídák’s multiple comparison test). (Q.) Total number of seizures during the recording period (p=0.0360, U-19.50). (R.) Average number of seizures per day (p=0.0551, U=21.50). (S.) Average duration of each seizure (p=0.0015, U=8.5). (T.) Cumulative seizure duration (p=0.0079, U=13.50). J-M: SE + Veh (n=11), SE + DQ (n=9); N-O: no SE (n=16), Se + Veh (n=17), SE + DQ (n=18). P: no SE (n=14), Se + Veh (n=16), SE + DQ (n=14). Q-T: + Veh (n=10), SE + DQ (n=9). J-M: Unpaired t-test; Q-T: Mann Whitney Test. Mean = +/- S.E.M.

As with p16-specific SC ablation, chronic DQ treatment after SE termination reduced p16 expression in the hippocampus (Fig. 4J) but did not affect microglial population numbers (Fig. 4K). Unlike with the genetic p16 ablation strategy, DQ-treatment reduced p16+ microglia both in the overall hippocampal cell population (Fig. 4L) and in the hippocampal microglia population (Fig. 4M). 8 weeks after SE, we assessed anxiety/exploratory behaviors (Supplementary Fig. 7A-D), recognition memory functions (Supplementary Fig. 7F-I), and spatial memory with (Fig. 4N-P). As with p16-specific SC ablation, senotherapy normalized SE-associated spatial memory impairments with a significant reduction in latency to escape in BM (Fig. 4O), and increased novelty preference index in NOLT (Fig. 4P). Moreover, DQ-treated mice had fewer seizures over 12 recording days (Fig. 4Q), while daily seizures did not differ from vehicle-treated mice (Fig. 4R). Senotherapy also reduced the average seizure duration (Fig. 4S) and cumulative seizure duration (Fig. 4T) These findings strongly suggest that continued intermittent DQ treatment following SE exerts an anti-epileptogenic effect and normalizes spatial memory impairments.

### Global microglial depletion throughout epileptogenesis does not impact seizure burden or associated cognitive impairments

The CSF1R receptor is a dependence receptor for microglia and is necessary for both microglial proliferation and survival.^58^ While several studies have published on the effects of inhibition of microglial proliferation and survival on seizures and epilepsy using CSF1R inhibitors to deplete microglia, reports in chronic epilepsy have been conflicting with some studies showing exacerbated seizures^59,60^ and others showing reduced seizures^61,62^.

Given that senescent microglia have exited the cell cycle, we hypothesized that the CSF1R antagonist, PLX3397 (PLX)^43,59,63,64^, would *spare* senescent microglia, and produce minimal impact on epilepsy and associated comorbidities. We depleted microglia with PLX immediately after SE termination (as with our senolytic studies), and continued treatment throughout the entirety of the study (4 months) to assess the impact on seizure burden and associated cognitive impairments, after which, brains were collected to confirm microglial depletion (Fig. 5A). PLX-treated SE mice had a 71% reduction in hippocampal microglia (Fig.5B-I), similar to what has been reported previously^59,63^. Total p16 (tdTom) hippocampal expression (Fig. 5K) and p16 expression in microglia remained unchanged, however, consistent with senescent-like microglia having exited the cell cycle and displaying resistance to apoptosis, there was a noticeable trend towards a higher proportion of p16+ microglia in PLX-treated mice (Fig. 5L).

**Fig. 5.**
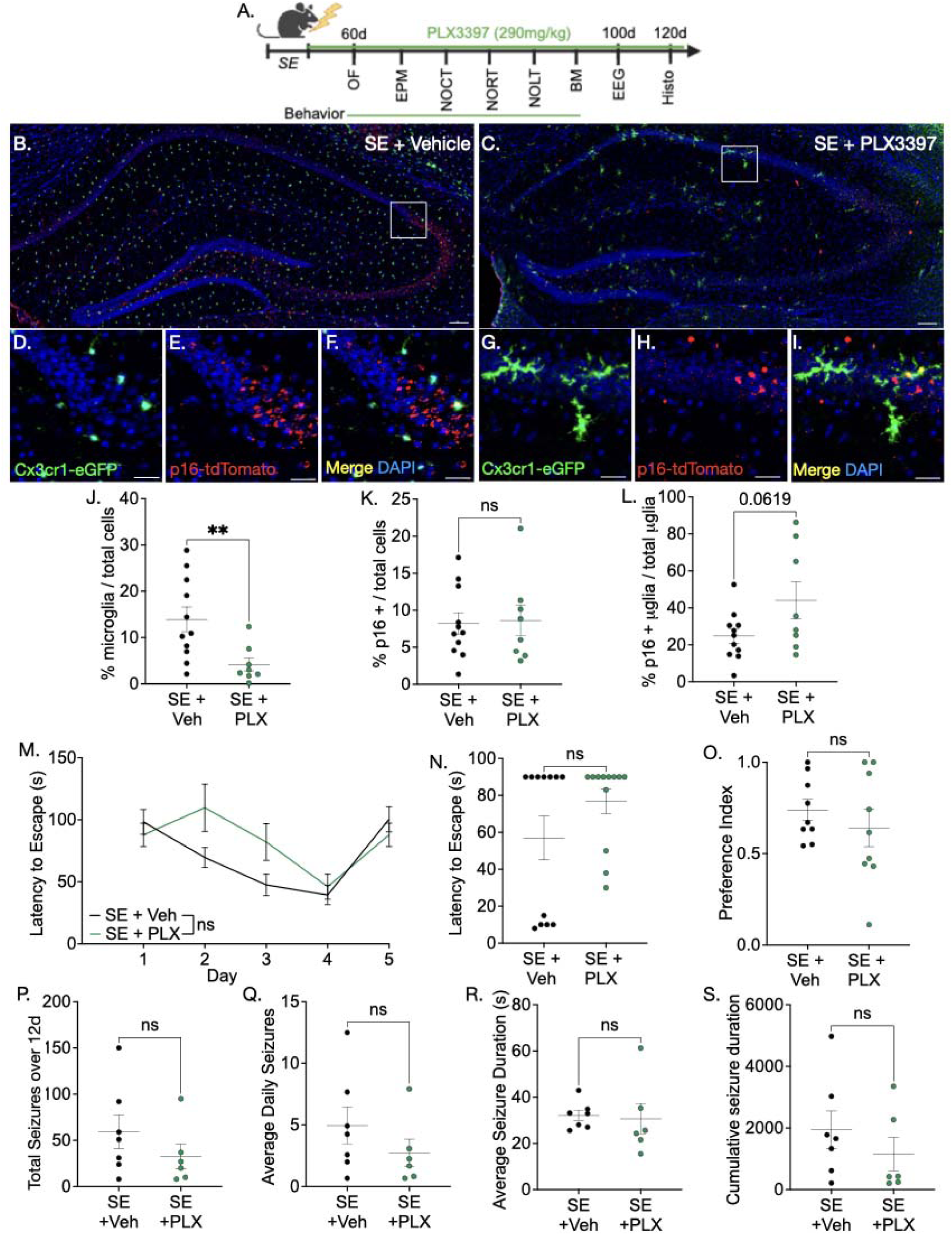
Global microglial depletion during epileptogenesis does not impact seizure burden or cognitive processes. (A.) Experimental timeline. 4-month-old male and female mice receive vehicle or PLX treatment following SE termination until the end of the experiment. 8 weeks after SE animals undergo behavioral tests, followed by 12 days of EEG implantation and EEG monitoring. At the end of the study, brains are processed for histological analysis of senescence markers. Representative image of p16-tdTom (red) and Cx3cr1 positive microglia (green) expression in (B.) vehicle-treated and (C.) PLX-treated mice 3 months following SE. Zoomed in representative image of p16, Iba1, and merged immunofluorescence in vehicle-treated (D-F.) and PLX-treated mice (G-I.). (J.) percent microglia from total detected nuclei (p=0.0068, U=12). (K.) percent tdTom positive cells from total detected nuclei (p=0.0203, U=16). (L.) percent tdTom positive microglia from total detected microglia (p=0.0619, t=1.999, df=17). (M.) Barnes maze training. p=0.104; t=1.64, DF=95.378; Mixed Model Analysis, Šídák’s multiple comparison test. Main effect trial day (p<0.0001, F(4,45.339)=10.961), no effect of treatment (p=0.104, F(1,95.378)=2.696) or interaction of day and treatment (p=0.085, F(4,45.339)=2.195). (N.) Barnes maze probe (p=0.3199, U=56). (O.) Novel Location preference index after a 5-minute delay (p=0.435, U=31). (P.) Total number of seizures during the recording period (p=3.112, U=13.50). (Q.) Average number of seizures per day (p=0.3112, U=13.5). (R.) Average duration of each seizure (p=0.3660, U=14). (S.) Cumulative seizure duration (p=0.3660, U=14). J-L: n=11 SE + Veh, n=8 SE + PLX. M-N: n=12 animals per group. O: n=9 animals per group. P-S: n=7 SE + Veh, n=6 SE + PLX. J-L; N-S: Mann Whitney Test. Mean = +/- S.E.M.

Unlike senolytic therapies, depletion of microglia (and predominantly non-senescent microglia) was without effect on behavioral tests assessing anxiety, exploration, recognition (Supplementary Fig. 8), and spatial memory functions (Fig. 5M-O) in SE mice. Similarly, we found no changes in total seizures over a 12-day recording period (Fig. 5P), average daily seizures (Fig. 5Q), average seizure duration (Fig. 5R), or cumulative seizure duration (Fig. 5S). These findings further solidify that senescence-like microglia contribute to epileptogenic processes as microglial depletion predominantly cleared non-senescent microglia while sparing senescent microglia.

### A p16-specific senescent phenotype in hippocampi from patients with TLE

Lastly, we investigated whether the robust senescence-like phenotype observed in our experimental TLE model is also observed in samples from patients with human TLE (hTLE). We obtained 9 autopsy control hippocampi and 11 resected hippocampi from patients with TLE from Georgetown University Brain Bank and assessed p16 expression and patient demographics. The average age of the patients with TLE at the time of seizure onset was 18 +/- 9.6 years with an average of 15 +/- 8.2 years with seizures before resection. The average age at resection was 33 +/- 13.3, whereas the average age of autopsy control patients was 65 +/- 12.8 years. While it is well established that p16 expression increases with age, we found higher p16 expression in the hippocampi from patients with TLE, despite being on average 32 years younger than the autopsy control tissue (Fig. 6A-C). We found an overall increase in p16 expression in all glial cell types in the hippocampi from patients with TLE compared to control: microglia (Fig. 6D), oligodendrocytes (Fig. 6E), astrocytes (Fig. 6F), but not neurons (Fig. 6G). Interestingly, we found an increased microglial density in hippocampi from patients with TLE (Fig. 6H), but not other cell types (Fig. 6I-K). We further examined the patient demographic data and did not find a significant correlation between p16 expression and age, years with epilepsy, or age of seizure onset (Table 1). There was a significant group difference between p16 expression and Engel class^65^, a clinical metric to measure the amount of seizure freedom following resection (p=0.0475, U=1, Mann-Whitney test, data not shown). These findings suggest that p16+ senescence-like phenotype is expressed predominantly in glia in TLE hippocampi and worse post-surgical outcome is associated with higher p16+ levels.

**Fig. 6.**
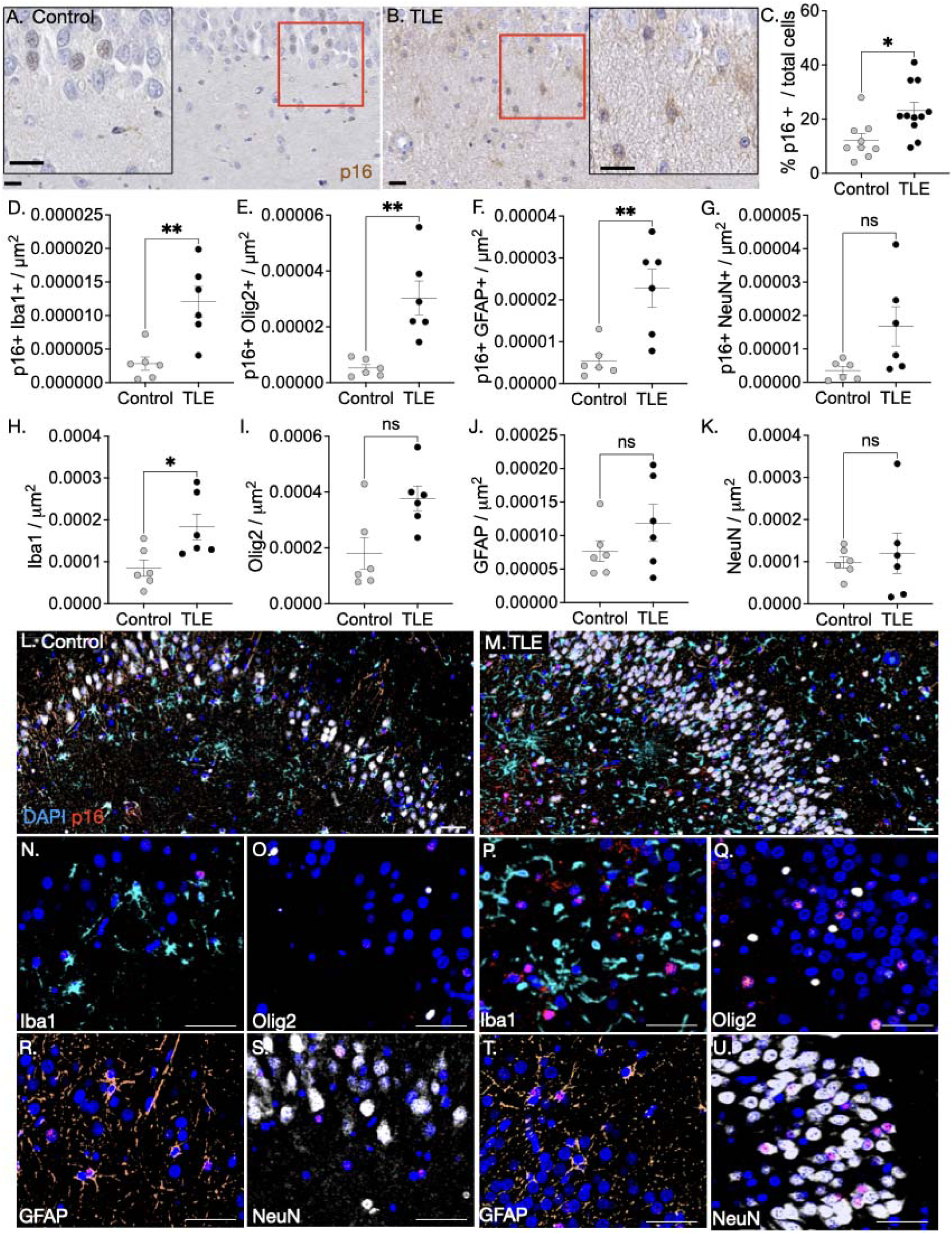
A p16+ senescence-like phenotype phenotype is prominent in resected hippocampi from patients with TLE. Representative photomicrograph of p16 expression in (A.) autopsy control hippocampus and (B.) resected hippocampus from patient with TLE. (C.) Quantification of percentage of p16+ cells in total detected nuclei in control hippocampi (n=9) and resected hippocampi from TLE patients (n=11) (p=0.0116, t=2.808, DF=18, Unpaired t-test). Density of p16 colocalization with cell types from control hippocampi and resected hippocampi from patients with TLE: (D.) microglia (p=0.0043, U=1); (E.) oligodendrocytes (p=0.0022, U=0); (F.) astrocytes (p=0.0087, U=2); (G.) neurons (p=0.0649, U=6). Cell type density in hippocampi of autopsy control and patients with TLE: (H.) microglia (p=0.026, U=4); (I.) oligodendrocytes (p=0.0649, U=6); (J.) astrocytes (p=0.3939, U=12), and (K.) neurons (p>0.999, U=18). Representative photomicrographs of p16 (red) multiplexed with microglia (Iba1, cyan), oligodendrocytes (Olig2, yellow), astrocytes (GFAP, orange), and neurons (NeuN, white) in (L.) autopsy control hippocampus and (M.) resected hippocampus from patient with TLE. Zoomed in representative images of p16 colocalization with (N.) Iba1, (O.) Olig2, (R.) GFAP, and (S.) NeuN in autopsy control hippocampus. Zoomed in representative images of p16 colocalization with (P.) Iba1, (Q.) Olig2, (T.) GFAP, and (U.) NeuN in hippocampus of patient with TLE. D-K: n=6 per group; Mann-Whitney Test. Mean = +/- S.E.M.

## Discussion

Here we demonstrate that hallmark indicators of cellular senescence accumulate in both the rodent and human brain during chronic epilepsy. Moreover, eliminating senescence-like cells after initiating SE reduces the development of epilepsy and associated comorbid behavioral deficits in experimental TLE. While previous studies have examined and targeted pathways that are downstream of senescence, (e.g., inflammation^12,17,66^, oxidative stress^67,68^, mTOR^69^, and p53^70–72)^, This is the first study to explicitly examine a senescence-like phenotype in the context of epileptogenesis.

A putative senescence phenotype was evident in our experimental TLE model 2 weeks after SE and continued to accumulate throughout the latent period and the period after the onset of spontaneous seizures. Similarly, increased numbers of p16+ cells were evident in resected hippocampi from patients with drug-resistant TLE. The difference in patient samples was robust, even in the face of substantial age differences between TLE and control groups (TLE cases were half the age of autopsy controls), suggesting that epileptogenic insults potently drive p16 expression. We found p16 expression across all glial subtypes (microglia, astrocytes, and oligodendrocytes) in hTLE samples, while microglia predominantly expressed p16 in our experimental TLE model.

Microglial senescence has been implicated in aging and neurodegenerative diseases such as Alzheimer’s disease (AD)^73,73,74^, which is increasingly recognized as a comorbidity of epilepsy^75–78^. We observed structural and functional alterations in microglia as a function of p16 expression in our epilepsy model. p16+ microglia displayed a shift in morphology towards a more activated state and reduced spontaneous process motility, indicating impairments in normal surveillance of their environment. Chemotaxic responses to purinergic stimulation, however, did not differ as a function of either epilepsy history or p16 expression. This differs from studies acutely following SE, in which enhanced microglial process velocity to a purinergic source has been reported^46,79–81^. Microglial responses to purines is concentration dependent; lower concentrations of ATP enable microglial response, whereas higher concentrations saturate ATP receptors and dampen the response^45^. Given reports of upregulated purinergic receptors in chronic epilepsy^82–85^, it is possible that we were at a saturating concentration. Together, the “activated” patterns in p16+ microglia may contribute to epileptogenesis.

Inflammatory processes involved in epilepsy overlap with components of the heterogeneous and complex SASP^17^. Interestingly, clearance of p16+ cells throughout epileptogenesis reduced the expression pro-inflammatory cytokines associated with an M1 microglial polarization state, such as IL-1β. This finding suggests that p16+ microglia are senescence-like in experimental TLE, as others have shown a senescence-independent phenotype in p16-expressing microglia is associated with anti-inflammatory (M2 polarized) states^86,87^.

We utilized two strategies to probe the contribution of senescence-like cells. We started senolytic treatment immediately after SE termination and continued treatment until the completion of the study (4 months of treatment) to achieve an anti-epileptogenic effect. We compared a genetic senolytic approach to a pharmacological senolytic approach. As an additional comparison, we evaluated the impact of CSF1R inhibition to reduce microglial proliferation and survival. As microglia predominantly express a senescence-like phenotype (and are thus resistant to cell death^24^) in our model, we expected that an inhibitor of microglial proliferation, PLX, would have a minimal impact on senescent microglia, but would selectively eliminate non-senescent microglia. Consistent with this, despite a 71% reduction in hippocampal microglia, neither p16 levels, cognitive function, nor seizure burden were impacted by CSF1R inhibition following SE. Another study demonstrated microglial depletion with PLX5622, another CSF1R inhibitor, reduced p16 expression in the aged mouse brain, however the study did not disclose the proportion of p16-positive vs. p16-negative microglia^88^. Our data suggest that senescence is the dominant phenotype; it is not microglial expansion and proliferation, but rather the escape of microglia into senescence, where they remain unaffected by the PLX inhibitor. Senolytic therapy also avoids systemic toxicity associated with global CSFR1 inhibition including anemia, leukopenia, and hepatotoxicity.^89^

In contrast to the inhibition of microglial proliferation, both genetic (INK-ATTAC) and pharmacological (DQ) senolytic interventions rescued spatial memory impairments and reduced seizure burden following epilepsy development. While spatial memory in the Barnes maze is clearly hippocampal dependent, other forms of memory (novel object recognition, object location memory) are not. While senolytic therapy improved hippocampal LTP, we did not examine if it rescued circuit dysfunction in other regions. It is likewise possible that residual memory deficits are due to damage and cell loss that are senescence-independent (e.g., a function of the initial excitotoxic SE injury). Rodent models of chronic epilepsy have reported worsened spatial memory functions more so than other types of memory processes^90^. We focused our observations exclusively on the hippocampus for this study, as it is a critical substrate for TLE. It is possible that p16 expression may differ in other limbic regions^91,92^.

While both genetic and pharmacological SC ablation strategies reduced p16 expression in the hippocampus by approximately 50%, p16+ microglia levels were decreased with DQ treatment but not in the case of INK-ATTAC. This level of senolysis was effective at extending healthspan and normalizing function in progeroid mice, atherosclerotic mice, metabolic disorders, and other age-associated disorders^33–35,54,55^. Moreover, we have previously found that this degree of senotherapy with DQ reduced age-related enhanced sensitivity to chemoconvulsants^38^. These strategies differ in the mechanism of action and may target different subsets of SCs. The INK-ATTAC mouse model eliminates cells that have high p16 expression^33^, while DQ disables the SCAPs that prevent SCs from apoptosis and does not specifically target p16 expressing cells^93^.

Despite the striking protection against spontaneous seizures, we observed (40% of mice in the ATTAC model were protected from spontaneous seizures), the mechanism(s) by which cellular senescence contributes to epileptogenesis remains unclear. Pathological changes associated with senescence and associated SASP include significant decline in neuronal survival, dendritic and axonal arborization, post-synaptic densities, dendritic spines, synapse, and cortical volumes^24^. Partial reduction in SASPs may allow glial cells to provide structural, metabolic, and trophic support to neurons, preserving normal hippocampal pathology and function, while still allowing glia to provide inflammatory signaling when necessary. Moreover, reduction in proinflammatory cytokine levels may reduce pathological excitation^9^.

Our experimental approach has some limitations, including the fact that the senolytic interventions target all SCs. While the senescence-like phenotype is predominantly expressed in microglia, other cell types do express a senescence-like phenotype, and they may also contribute to epileptogenesis. For example, senescent neuroblasts have been implicated in an age-associated decline in neurogenesis,^94^ and we detected p16+ doublecortin positive cells in the present study. Similarly, senescent astrocytes have been reported in aged human hippocampi^95^. We detected colocalization of p16+ and GFAP in both tissue from human TLE and in post-SE mice; astrocyte dysfunction has likewise been proposed as a contributor to epileptogenesis^62,85,96,97^. Finally, oligodendrocyte precursor cell senescence was shown to drive cognitive decline in a mouse model of amyloid-beta pathology^27^, and again we detected colocalization between p16 and Olig2 in both human and mouse tissue. Oligodendrocyte dysfunction has been proposed to contribute to epileptogenesis through maladaptive myelination^98^. Collectively, these studies point towards a diverse and multifaceted role of senescence in the brain, manifesting across distinct cell types.

Ultimately, these findings point towards an involvement of senescence-like cells in seizures and the development of epilepsy after injury. Given that *all* current therapies for the epilepsies are symptomatic, i.e., they reduce seizures, but do not prevent the development of epilepsy, the striking protection against epileptogenesis and associated cognitive comorbidities we describe with senotherapy suggests a novel mechanism for therapeutic development in the treatment of epilepsies.

## Supporting information

Supplemental Results

## Illustrations

All illustrations were created with BioRender.com.

## Data Availability

Raw data are available upon request.

## Code Availability

Microglia morphology analysis available on https://github.com/forcelli-lab/3DMorph_HPC_OPT

## Author Contribution

Conceptualization: T.K. and P.A.F.; Data curation: T.K. and P.A.F.; Formal analysis: T.K., D.J.M., A.I.H., L.A.F., T.P.C., M.C.S., I.S., N.M.K., M.C., J.A.B. and S.V.; Funding acquisition: T.K. and P.A.F.; Investigation: T.K., D.J.M., A.I.H., L.A.F., T.P.C., M.C.S., I.S., N.M.K., M.C. and J.A.B.; Methodology: N.M.K., B.T.H. and S.V.; Project administration: T.K. and P.A.F.; Software: N.M.K.; Supervision: B.T.H., S.V. and P.A.F.; Visualization: T.K., D.J.M. and P.A.F.; Writing – original draft: T.K. and P.A.F.; Writing - review & editing: T.K., D.J.M., A.I.H., L.A.F., T.P.C., M.C.S., I.S., N.M.K., M.C., J.A.B., B.T.H., S.V. and P.A.F.;

## Acknowledgements

This work was supported by R21NS125552 to P.A.F. T.K. was supported by F99NS129108 and T32NS041218. D.M. was supported by T32GM142520. A.I.H. was supported by T32GM144880.

## Methods

### Animals

All experiments and procedures were carried out in accordance with the IACUC policy at Georgetown University Medical Center (Protocol # 2019-0037). Mice were housed at 22 ℃ with a 12 h light/dark cycle from 6am to 6pm, and allowed for free access to food and water. Unless otherwise indicated, 12 to 16 week old male and female mice were used.

### C57BL/6J

Wildtype C57BL/6J mice (RRID:MGI:3028467) were obtained from Jackson Laboratory (Bar Harbor, ME, USA), for breeding purposes and senolytic studies.

### p16^tdTom^ Reporter

For senescence characterization studies, p16 tdtomato reporter mice were used. This reporter strain was generated by Norman Sharpless’s group, in which tandem-dimer Tomato (tdTom) fluorochrome was “knocked in” into exon 1a of the p16^INK4a^ locus, and allows for the visualization, enumeration, isolation, and characterization individual p16^INK4a^ expressing cells ^41^. Thus, this strain retains the p16^INK4a^ allele, with tdTom as the second allele, allowing for p16^INK4a^ promoter activation to be measured. tdTom positive mice were maintained on a C57BL/6J background.

### CX3CR1^eGFP/+^ Reporter

CX3CR1^GFP/+^ (RRID:IMSR_JAX:005582) mice express green fluorescent protein (GFP) in place of one allele of the fractalkine receptor gene, resulting in fluorescently labeled microglia ^50^. Mice were previously backcrossed to a C57BL/6J congenic background.

### INK-ATTAC

For genetic senescent cell ablation studies, INK-ATTAC mice were used. INK-ATTAC (p16**^INK^**^4a^ **A**poptosis **T**hrough **T**argeted **A**ctivation of **C**aspase) line was created by James Kirkland’s group at the Mayo Clinic to examine p16 specific depletion of senescent cells ^33^. The p16^INK4a^ promoter harbors a suicide gene, a membrane-bound myristoylated FK506-binding-protein–caspase 8 (FKBP–Casp8) fusion protein expressed specifically in p16^INK4a^ positive cells, which becomes activated by an exogenous synthetic dimerizer, AP20187, and selectively kills p16^INK4a^ positive senescent cells. These mice were on a 129 scv C57BL/6 FVB mixed background.

### Status Epilepticus

We used the pilocarpine model of epilepsy (Turski et al., 1984). Pilocarpine is a chemoconvulsant that is used to induce Status Epilepticus (SE), in which prolonged seizures cannot self-terminate. Animals were pretreated with scopolamine methyl bromide (Sigma-Aldrich, S8502) and terbutaline hemi sulfate (Sigma Aldrich, T2528) (both i.p., 2mg/kg) to block peripheral effects of pilocarpine and dilate the respiratory tract, as conducted in Cho et al 2015. 30 minutes after, 280mg/kg pilocarpine hydrochloride (Cayman Chemicals, c23131841) was administered intraperitoneally. Acute seizures were behaviorally monitored using the following seizure rating scale (stage 1, facial clonus; stage 2, unilateral forelimb clonus; stage 3, bilateral forelimb clonus; stage 4, rearing with forelimb clonus; stage 5, rearing and falling with forelimb clonus; stage 6, jumping and running). SE was defined by 5 minutes of stage 3 continuous tonic clonic convulsive seizures. Mice were monitored for two hours until seizure activity was reduced with diazepam (i.p., 5 mg/kg; Dash Pharmaceuticals). After termination of SE, mice were administered 5% dextrose solution to facilitate their recovery and improve survival.

### Treatment

Animals received the respective treatment immediately after termination of SE. C57BL/6 mice were randomized to receive pretreatment of senolytic cocktail, Dasatinib (i.p., 5mg/kg, Selleckchem, S1021) quercetin (i.p, 50mg/kg, Sigma-Aldrich, Q4951) or vehicle once a week until the end of the experiment. INK-ATTAC mice were randomly assigned to receive FKBP-homodimerizer, AP20197 (i.p, 2mg/kg, MedChem Express, HY-13992).

### Surgery

For seizure detection, mice were implanted with wireless EEG telemeters (DSI-HDX02, EMKA easyTEL S) for chronic seizure monitoring. This method reduces stress compared to tethered systems and allows mice to freely roam in their home cage ^99,100^. Animals were anesthetized and placed into a stereotaxic frame using blunt earbars. Before any incisions, bupivicaine was administered subcutaneously (s.c.) to the dorsal back of the mice. Carprofen was administered (s.c.) to the scalp before the skin of the scalp is incised (midline incision ∼1cm) and retracted. Bregma was marked and two holes were bore through the skull using a dremel tool, sterile saline or PBS was added at the craniotomy site to reduce heating. Four stainless steel epidural cortical electrodes insulated except at the tip were implanted bilaterally over the parietal cortex. This configuration has no common lead, such that each pair of leads is coupled into an instrumentation amplifier (i.e.; differential inputs). Straight hemostats were used to create a subcutaneous pocket posterior to the head and thread the telemeter into the pocket. The leads were fixed to rest on top of the dura in a small groove made in the skull using cyanoacrylicate gel. The skin was sutured using 3-0 or 4-0 absorbable monofilament and a simple interrupted buried suture pattern. Vet bond was placed over the suture line as a secondary form of closure. The wound margin was treated with lidocaine ointment and triple antibiotic. Animals were placed on a heating pad to recover and given a 1ml injection of normal saline (s.c.). Animals were single housed for EEG monitoring following surgery to allow for individual seizure detection.

### EEG monitoring

Animals were placed in their individual cages which were placed on an MX2 data exchange matrix receiver for simultaneous video-EEG recordings. Mice were recorded for 12 days.

### EEG analysis

EEG signals were bandpass filtered (1–50 Hz) and manually scored using Labchart v8 1.28 software. For each animal, we calculated the (1) daily number of seizures, which was defined as the count of seizures during a 12 day period; (2) the average seizure duration, which was defined as the average of the lengths of individual seizures during the 12 day period; (3) the total seizure burden, which was the total duration of electrographic seizure activity, and (4) cumulative seizure duration, which was the average duration of seizures multiplied by the total number of seizures in the recording period; reflecting a combined metric of both seizure number and seizure duration.

### Behavior

Two months following SE, mice underwent a battery of behavioral tests, described below. Any-maze software was used for video tracking. For recognition tests: exploration of an object was defined as the animals’ nose being within 1 cm of and oriented toward the object, sniffing at, or otherwise closely attending to the object. Behavior analyses were conducted manually, observers remained blinded to experimental conditions.

### Open Field

Locomotion activity was monitored in an open field box. Mice were habituated for one hour. All novel recognition tests were performed in the same apparatus, a 16“wL×L16“dL×L16“h box with a wooden base and clear glass plexiglass containers.

### Elevated Plus Maze

Mice are placed at the junction of the four arms of the maze, facing an open arm, and entries/duration in each arm are 3 min. The maze is elevated (height).

### Novel Context Recognition Test

Two distinct chambers were used for this test, a covered circular chamber with bedding, containing 2 red cylinders; and a square open field chamber containing 2 blue cubes, the same size as the cylinders. Animals habituated in the chamber for 2 days without objects; first 10 minutes in the circular chamber, immediately followed by 10 minutes in the square chamber. On day 3, probe day, animals were habituated with the respective chambers with objects—5 minutes in the circular chamber, followed by 5 minutes in the square chamber. After a 5 minute delay, the mice were placed back in the circular chamber in which one of the red cylindrical objects were replaced by a blue cube object (associated with the square chamber). During this test, mice were given 3 minutes to explore the objects.

### Novel Object Recognition Test

Animals were familiarized in the open field box with two objects (pumpkins) for five minutes. Following a delay of 5 minutes or 24 hours in an empty cage, mice were placed back in the testing chamber, with one of the pumpkins replaced with a yellow toy duck of the same size. Animals were allowed to explore the objects for 3 minutes during probe phases.

### Novel Location Recognition Test

Animals were familiarized in the open field box with two objects (silver pyramids) for five minutes. Following a delay of 5 minutes or 24 hours in an empty cage, mice were placed back in the testing chamber, with one of the silver pyramids moved to a new location. Animals were allowed to explore the objects for 3 minutes during probe phases.

### Barnes Maze

The Barnes Maze is used to study spatial memory and navigation. The maze consists of an elevated circular platform (92cm) with 20 holes (5cm) around the perimeter (Maze Engineers). The target hole consisted of an escape box, and the remaining holes had a false floor. The location of the escape box remained consistent, and animals used spatial cues to find the escape platform. Aversive overhead lighting and radio static were used to motivate the mice to find the escape platform. Animals were trained for 4 consecutive days, 4 trials each day with an inter-trial interval of 15 minutes. Each trial was 180 seconds long. If the animal failed to find the target hole, it was guided to the escape hole. Animals were kept in the escape box for 30 seconds. 24 hours following the fourth training day, animals were subjected to the probe trial where they were given 90 seconds to find the escape platform. The chamber was cleaned thoroughly between animals. Latency to platform was used to measure learning each day during training and for probe.

### Acute Slice Preparation

Mice were anesthetized with unmetered isoflurane (Patterson Veterinary) and intracardially perfused in ice-cold sucrose N-Methyl-D-glucamine (NMDG) solution containing NMDG 92 mM, KCl 2.5 mM, NaH2PO4 dihydrate 1.25 mM, NaHCO3 30 mM, HEPES 20 mM, glucose 25 mM, sucrose 10 mM, ascorbic acid 5 mM, thiourea 2 mM, sodium pyruvate 3 mM, N-acetyl-L-cysteine 5 mM, MgSO4 heptahydrate 10 mM, and CaCl2 dihydrate 0.5 mM at pH 7.3–7.4 and osmolarity 300–310 mOm/kg. 300 μm horizontal hippocampal slices were cut on a Vibratome 3000 Plus Sectioning System (Vibratome, St. Louis, MO, USA), in ice-cold NMDG. Sections were recovered NMDG for 5 minutes 32 °C and then transferred to a modified artificial cerebrospinal fluid (aCSF) solution for 30 minutes at 32°C, containing NaCl 120 mM, KCl 3.5 mM, NaH2PO4 1.25 mM, NaHCO3 26 mM, CaCl2 dihydrate 1 mM, MgCl2 7 mM, and dextrose 10 mM at pH 7.3–7.4 and osmolarity 300–310 mOm/kg. Slices were transferred to room temperature (22–24 °C) and equilibrated for > 10 min before use. Slices were then transferred to room temperature (RT = 22-24 °C) and maintained in recording aCSF solution containing in mmol/L: 124 NaCl, 3.5 KCl, 1.2 Na2HPO4, 26 NaHCO3, 2 CaCl2, 1 MgCl2 and 10 dextrose at a pH of 7.3-7.4 and osmolarity 300-310 mOm/kg. To limit artifactual microglial activation from dissection/sectioning, all slices were used within 3 hours of euthanasia and all microglia studied had somata at least 30 μm away from the cut surfaces. aCSF solutions were maintained at pH = 7.4 by bubbling with carbogen gas (95% O2/5% CO2, Roberts Oxygen, Rockville, MD, USA). All experiments were conducted at RT.

### Confocal Imaging

Confocal Z- and ZT-stacks were taken with a laser scanning microscope system (Thor Imaging System Division) equipped with 488/561/642 nm laser and Green/Red/Far-red filters and mounted on an upright Elipse FN1 microscope (Nikon Instruments). 284 × 284 x 20 μm (xyz) volumes of hippocampal slices were imaged through the 60x water-dipping objective (CFI Fluor 60XW, NA=1.0, WD= 2mm, Nikon). Differential interference contrast images were used to identify region of interest. 1024×1024 pixel ZT-stack images from 10 planes 1.5 μm apart were taken every 30 seconds. Baseline motility was imaged for 30 minutes in DG, CA3, and CA1 hippocampal subregions. For reactive microglia motility, a patch pipette containing 3mM ATP in aCSF was lowered 30 μm into the tissue to allow ATP passive diffusion into the tissue without pressure in the pipette. The surrounding volume was imaged for 30 minutes.

### Microglial motility analysis

Time-lapse images were manually cropped, then thresholded and binarized in the region of interest (ROI). Binary ROI from each timepoint t(x) were color-merged with a t(x+1) ROI, generating images containing red pixels (retraction), green pixels (extension), and yellow pixels (static). The numbers of each colored pixel were counted using the Color Counter plugin (Wayne Rasband in FIJI). Motility index was calculated for each pair of frames (t(x) and t(x+1)) such that measured extension and retraction over static. The motility index of each paired frame (e.g., t0+t1, t1+2, etc) was averaged for the whole 30 minute time-lapse.

### Microglia process velocity analysis

Microglia process velocity was quantified manually in FIJI using the Manual Tracking plugin (Fabrice Cordelieres). 3-5 processes were manually tracked over time per microglia that were moving towards the ATP containing pipette. 3-5 slices were analyzed per animal. Experimenters remained blinded to p16 expression in microglia while analyzing.

### Automated Assessment of Microglia Morphology

Morphology of microglia was assessed using confocal z-stacks. Iba1 stained cells were imaged at 0.0749 µm per x-y pixel and 0.0749 µm per z-step. The field of view was 2984×2500 pixels with 70-120 z-steps per image. Three regions, CA1, CA3, and DG, were imaged across three slices per animal. Images were pre-processed in ImageJ/FIJI to remove out-of-focus z-layers and separate channels. This was done to reduce memory needs for processing. The assessment parameters for each image were generated using 3DMorph Interactive mode (https://github.com/ElisaYork/3DMorph, Accessed December 1, 2023) to generate the noise reduction, threshold, and segmentation parameters for processing. 3DMorph is a MATLAB based script for analysis of microglia morphology in three dimensions. [York et al., 2018] The parameters generated in Interactive mode were fed into a modified version of 3DMorph, 3DMorph_HPC_OPT. 3DMorph_HPC_OPT is modified for optimization using a Slurm-based high-performance computing center. Modifications included optimization to the algorithms used to determine branch length, changes made to memory allocation processes, and output files. The automated assessment generated cell volume, cell territory (the convex hull encapsulating the cell representing the area surveilled by the cell), ramification index (territorial volume divided by cell volume), branch lengths, and number of endpoints. The presence of p16 was determined by user assessment of max projection images.

### Long Term Potentiation

Animals were perfused with NMDG. Brains were cut into 275 um-thick sections, incubated in NMDG at 32℃ for 30min, then a HEPES bath at room temperature for 4hrs. Slices were recorded in aCSF at 32℃. Slices were visualized using an upright microscope (Axioscope 2 FS Plus, Zeiss, Germany) with a 2.5x Plan-NEOFLUAR objective and camera. Recordings were filtered with a 1kHz high-pass filter and digitized at 20kHz using a MultiClamp 700B amplifier. Electrodes for recording potentials were placed in the stratum radiatum of the CA1 of the hippocampus. The stimulating electrode was placed near the Schaffer collaterals. Input/output curves for fEPSPs were recorded, and a stimulation intensity was selected from the curve that produced ∼30-50% of a maximum response. Slices were stimulated every 30sec for 10min to ensure a stable baseline signal. Slices were tetanized using theta burst stimulation (12 trains of 4 pulses (50us) delivered at 100Hz), as described in (Abrahamsson et al. 2016). Following tetanization, fEPSPs were recorded for 1hr. Recordings were analyzed using Clampfit v11.1.

### Protein Extraction and ELISA

Animals were perfused with ice cold PBS and their hippocampi were dissected and frozen with RIPA buffer with protease and phosphatase inhibitor. Hippocampi were sonicated on a sonicator (info) and spun down at 13K RPM for 30 minutes at 4C. The supernatant was collected. BCA was run (Pierce BCA kit, Thermofisher details) for protein concentrations. The samples were undiluted for ELISAs. A cytokine release multiplex ELISA panel (Biolegend) was done following manufacturer’s instructions.

### Immunohistochemistry

Mice were anesthetized and perfused transcardially with 1x PBS followed by 4% paraformaldehyde (PFA) in PBS. Brains were removed and postfixed in 4% PFA overnight, then cryoprotected in 30% sucrose in 0.1LM PBS. Brains were flash frozen and sectioned at 30um thickness on a cryostat (Leica CM 1850).

For staining, slides were washed with 1x PBS with 0.1% Triton-× 100 3 times for 5 minutes each. Nonspecific binding was blocked with Animal-Free Blocker and Diluent (Vector Laboratories, SP-5035-100) for 1 hour at room temperature. Antibodies were incubated overnight at 4°C [mouse anti-p16, monoclonal (Abcam Cat# ab54210, RRID:AB_881819; 1:500); mouse anti-human p16 (BD Biosciences Cat# 550834, RRID:AB_2078446; 1:500); mouse anti-p21 (Santa Cruz Biotechnology Cat# sc-6246, RRID:AB_628073;1:500); goat anti-tdTomato (SICGEN Cat# AB8181, RRID:AB_2722750; 1:500); rabbit anti-Iba1 (FUJIFILM Wako Pure Chemical Corporation Cat# 019-19741, RRID:AB_839504; 1:1000); rabbit anti-GFAP (Agilent Cat# Z0334, RRID:AB_10013382; 1:1000); rabbit anti-olig2 (Sigma-Aldrich Cat# AB9610, RRID:AB_570666; 1:500); guinea pig anti-doublecortin (Millipore Cat# AB2253, RRID:AB_1586992; 1:2000)]. On the second day primary was washed off in 3 washes in PBS. The slides were incubated with Alexa Fluor conjugated secondaries at 1:1000 (Jackson ImmunoResearch Laboratories Inc) for 1 hour at room temperature. The tissue was washed 3 times for 5 minutes each with PBS, quenched for autofluorescence (Vector Laboratories, SP8400-15), and cover slipped with VECTASHIELD Vibrance® Antifade Mounting Medium with DAPI (Vector Laboratories, H-1800). Sections were observed by fluorescence microscopy.

### SPiDER-β-gal

For the SPiDER-β-gal stain, tissue sections were incubated in 20 µM SPiDER-β-gal (Dojindo, SG02-10) in solution in 0.1% Triton in PBS for 60 min at 37°C. After washing of tissue sections, nuclei were stained with DAPI, and tissue sections were mounted with VECTASHIELD Vibrance® Antifade Mounting Medium with DAPI (Vector Laboratories, H-1800). Sections were observed by fluorescence microscopy (Leica Mica).

### RNAscope

Mouse brains were fixed overnight in 10% NBF and transferred to 70% EtOH. Brains were sectioned at 5 µm on a rotary microtome and stored at RT until use in RNAScope assay. Slides were baked at 60°C for 1 hr and then deparaffinized in xylene 2×5’ at RT and dehydrated with 100% EtOH 2×2’ and dried for 5’ at 60°C. The RNAScope assay proceeded according to the manufacturer’s protocol, following the guidelines for standard tissue processing of FFPE tissue. The RNAscope™ Multiplex Fluorescent V2 Assay was performed by Georgetown University’s Histopathology Tissue Shared Resource (HTSR). The probes used in this study were: p16 (Mm-Cdkn2a-tv2 44749), p21 (Mm-Cdkn1a 40855), Iba1 (Mm-Aif1 31914) and Il6 (Mm-Il6 315899). Staining was performed on in accordance to manufacturers’ instructions by HTSR. Tissue was analyzed in QuPath by thresholding object classifier.

### Human Tissue

We received 11 resected hippocampi from TLE patients and 9 autopsy control hippocampi from Georgetown University Brain Bank (IRB Study ID: STUDY00004786). Sections of 5μm thickness were cut from FFPE tissue blocks containing autopsy control and TLE hippocampal samples. The slides were baked at 60°, deparaffinized in xylene, rehydrated, washed in tap water and incubated with 10% neutral buffered formalin (NBF) for an additional 20 minutes to increase tissue-slide retention. Epitope retrieval/microwave treatment (MWT) for all antibodies was performed by boiling slides in Antigen Retrieval buffer 6 (AR6 pH6; Akoya, AR6001KT). Protein blocking was performed using antibody diluent/blocking buffer (Akoya, ARD1001EA) for 10 minutes at room temperature. The staining was performed using Leica Bond autostainer counterstained with spectral DAPI (Akoya FP1490) for 5 min and mounted with ProLong Diamond Antifade (ThermoFisher, P36961) using StatLab #1 coverslips (CV102450). The order of antibody staining and the antibody/OPAL pairing was predetermined using general guidelines and the particular biology of the panel. General guidelines include spectrally separating co-localizing markers and separating spectrally adjacent dyes. Multiplex IHC was optimized by first performing singleplex IHC with the chosen antibody/OPAL dye pair to optimize signal intensity values and proper cellular expression, followed by optimizing the full multiplex assay.

### Human Tissue imaging and analysis

Slides were scanned at 10X magnification using the Phenocycler Fusion system Automated Quantitative Pathology Imaging System (Akoya Biosciences). Whole slide scans were viewed with Phenochart (Akoya) which also allows for the selection of high-powered images at 20X (resolution of 0.5mm per pixel) for multispectral image capture. Multipspectral images of each tissue specimen were captured in their entirety. Multispectral images were unmixed using a spectral library built from images of single stained control tissues for each OPAL dye using the inForm Advanced Image Analysis software (inForm 2.8; Akoya). A selection of 10-15 representative multispectral images spanning all tissue sections was used to train the Qupath software (tissue segmentation, cell segmentation and phenotyping tools). All the settings applied to the training images were saved within an algorithm to allow batch analysis of all the multispectral images for the project. Nuclei were reacted with DAPI (AKOYA Biosciences, FP1490). For p16 chromogenic image analysis, nuclei were detected with a modified Stardist script in Qupath. Two observers trained a detection classifier for each image by selecting positive and negative p16 cells within heterogeneous regions of each image.

### Statistical Analysis

All data are expressed as mean ± S.E.M. Experimental groups were assigned by simple randomization. Data were collected blind. Graphpad prism (version 10.0.2) software and SPSS (version 29) were used for statistical analysis. Unless otherwise indicated, data were not evaluated for normality or heteroscedasticity.

## Extended Data Figures

**Supplemental Table 1.**
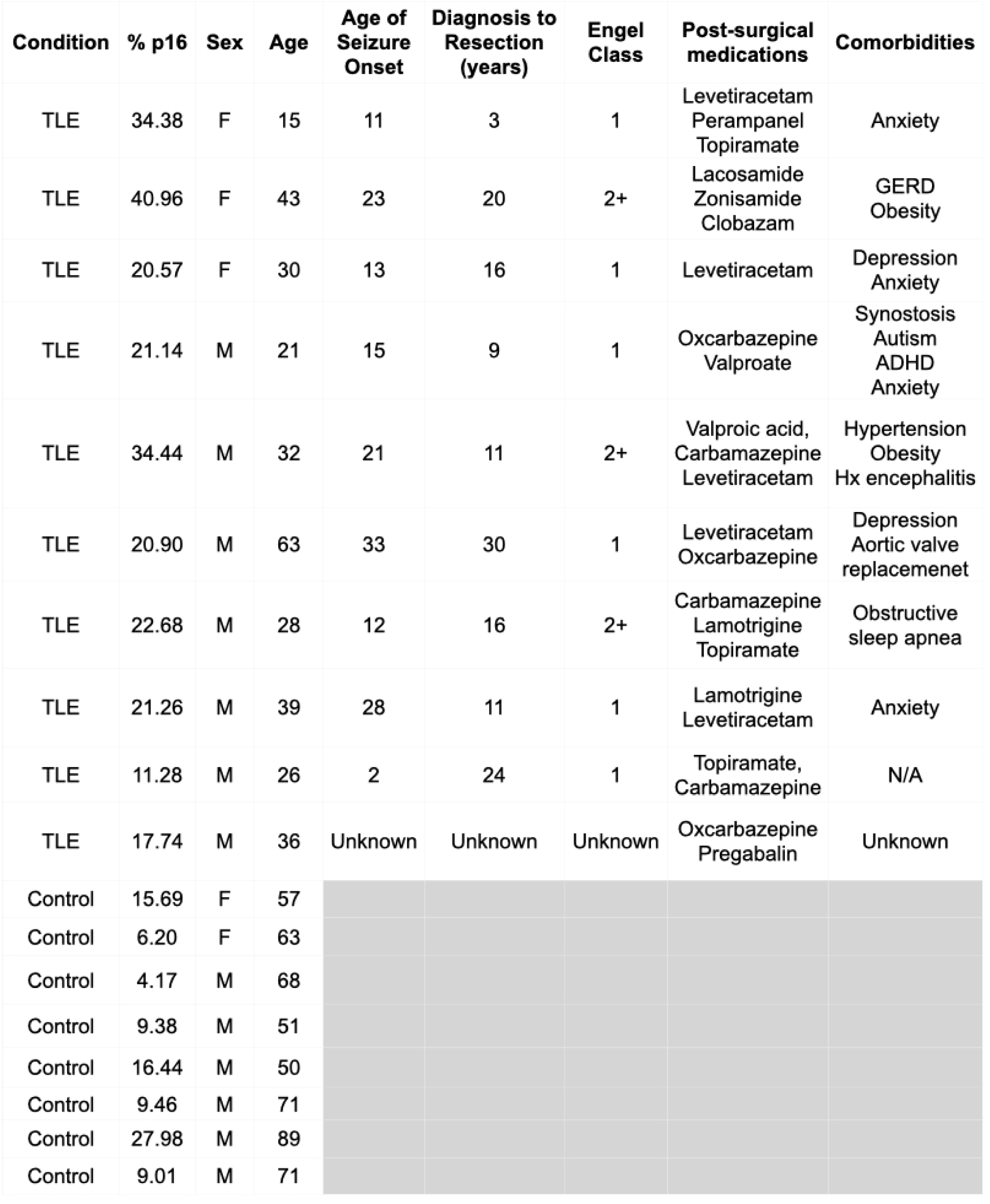
Demographics of patients with TLE and autopsy controls.

**Supplementary Fig. 1.**
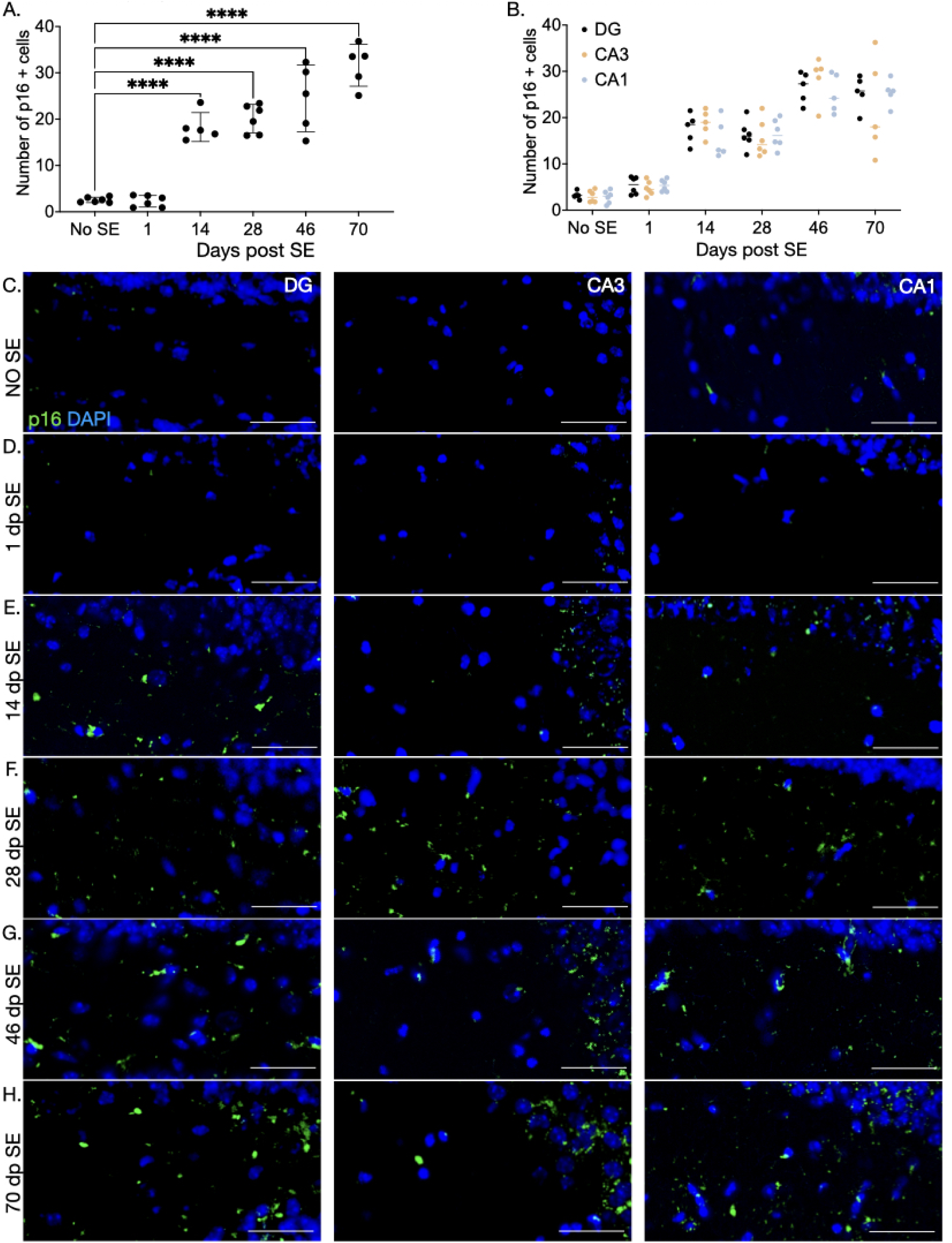
Immunofluorescence time course of p16 expression after SE. (A.) Quantification of p16 antibody at 1, 14, 28, 46, and 70 days post SE (p<0.0001, F(5,27)=54.78, One-Way ANOVA, Holms-Šídák’s multiple comparison test). No SE vs. 1 day: p=0.8978, t=1.297, DF=27; No SE vs. 14 days: p<0.0001, t=6.868, DF=27; No SE vs. 28 days: p<0.0001, t=8.029, DF=27; No SE vs. 46 days: p<0.0001, t=9.565, DF=27; No SE vs. 70 days: p<0.0001, t=12.69, DF=27. One-Way ANOVA, Holms-Šídák’s multiple comparison test. (B.) p16 antibody immunolabeling in hippocampal subregions: dentate gyrus (DG), CA3, and CA1 at 1, 14, 28, 46, and 70 days post SE. p=0.6158, F(1.618, 43.69)= 0.4221(Two-Way ANOVA, Holms-Šídák’s multiple comparison test). Main effect of days post SE (p<0.0001, F(5,27)=61.62), animal (p=0.0001, F(27,54)=3.275), no main effect of region (p=0.6158, F(1.618, 43.69)=0.4221); no interaction effect of days post SE x region (p=0.2954, F(10,54)=1.227). Representative images of p16 expression in DG, CA3, and CA1 in (C.) No SE, (D.) 1 day post SE, (E.) 14 days post SE, (F.) 28 days post SE, (G.) 46 days post SE, and (H.) 70 days post SE. Mean = +/- S.E.M.

**Supplementary Fig. 2.**
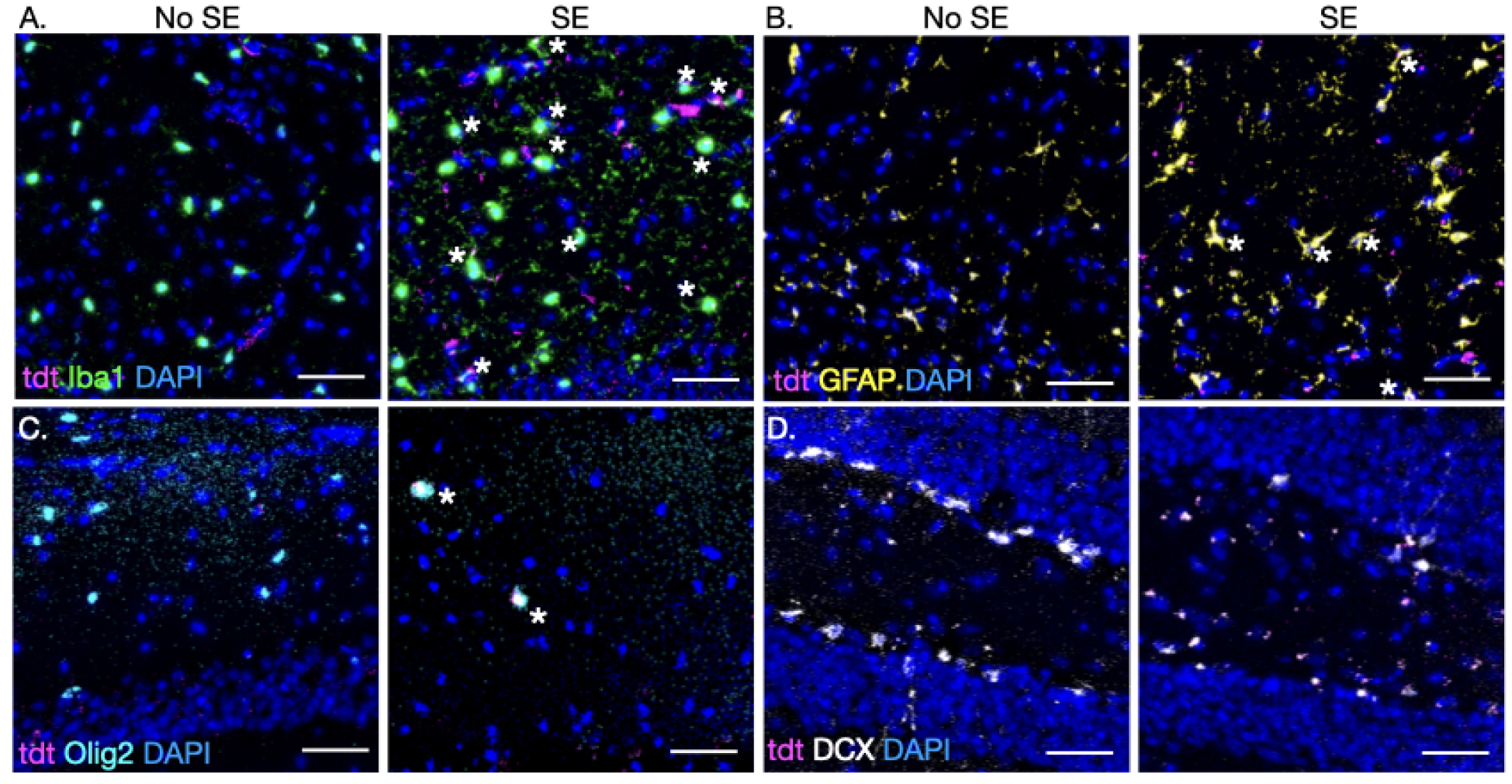
p16-tdTom reporter colocalization with cell-type specific markers. Representative photomicrographs of p16tdTom colocalization in non-seizure control (No SE) and chronic epilepsy (SE) mice immunolabeled with (A.) Iba1 for microglia, (B.) GFAP for astrocytes, (C.) Olig2 for oligodendrocytes, and (D.) Doublecortin (DCX) for immature neurons. Astericks denote colocalization.

**Supplementary Fig. 3.**
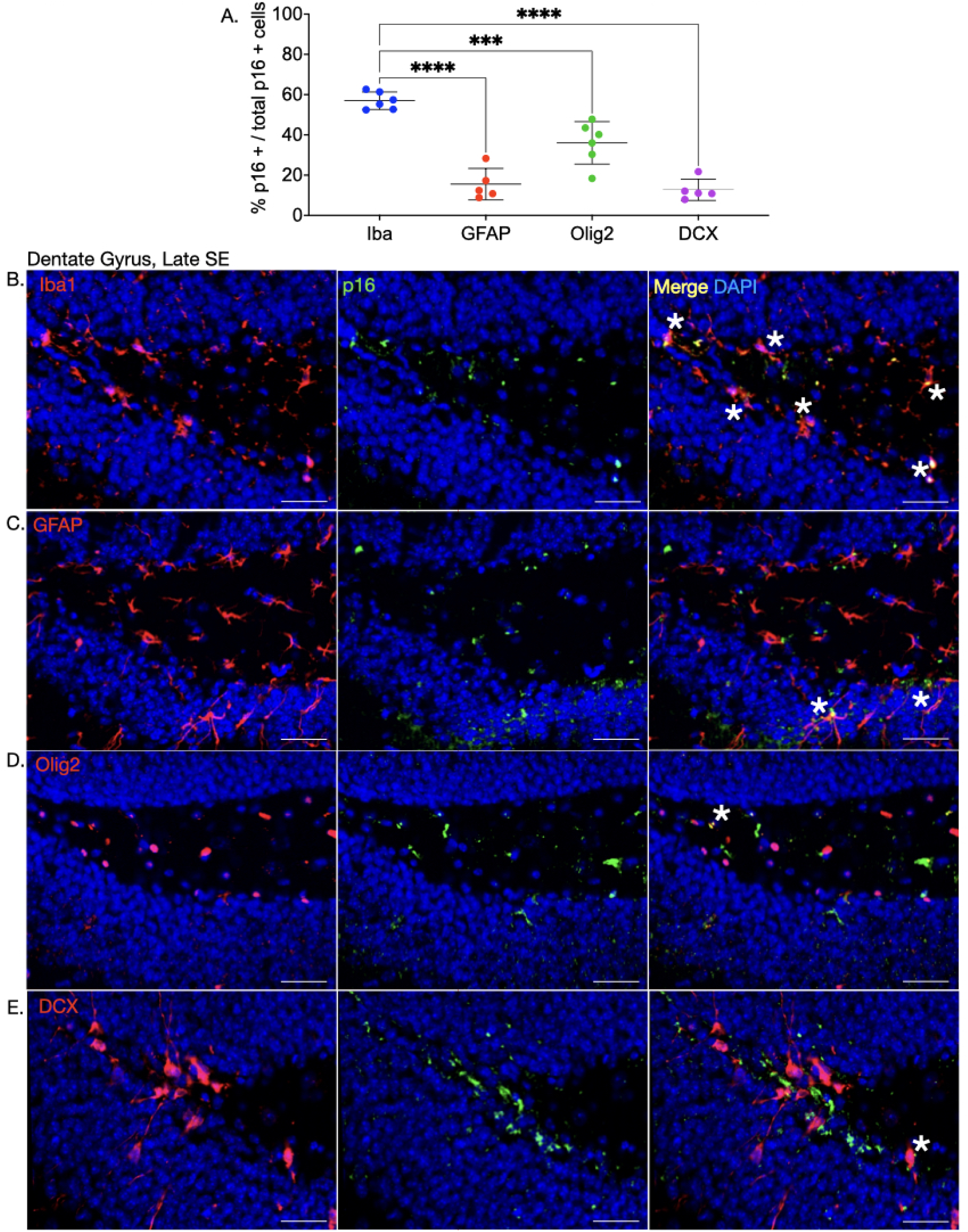
Immunofluorescence colocalization of p16 antibody and cell-type specific markers. (A.) Quantification of p16 antibody colocalization with mitotic cell markers: Microglia (Iba1, blue), Astrocytes (GFAP, red), Oligodendrocytes (Olig2, green), Immature Neurons (DCX, magenta). p<0.0001, F(3,18)=42.08; One-Way ANOVA, Holms-Šídák’s multiple comparison test. Iba1 vs. GFAP (p<0.0001, t=9.155, DF=18), Iba1 vs. Olig2 (p=0.0001, t=4.856, DF=18), Iba1 vs. DCX (p<0.0001, t=9.779, DF=18). Mean = +/- S.E.M. Representative photomicrographs of and cell type colocalization (red, left panel), p16 (green, middle panel), and merge (right panel) in the dentate gyrus of (B.) Iba1, (C.) GFAP, (D.) Olig2, and (E.) DCX. Astericks denote colocalization.

**Supplementary Fig. 4.**
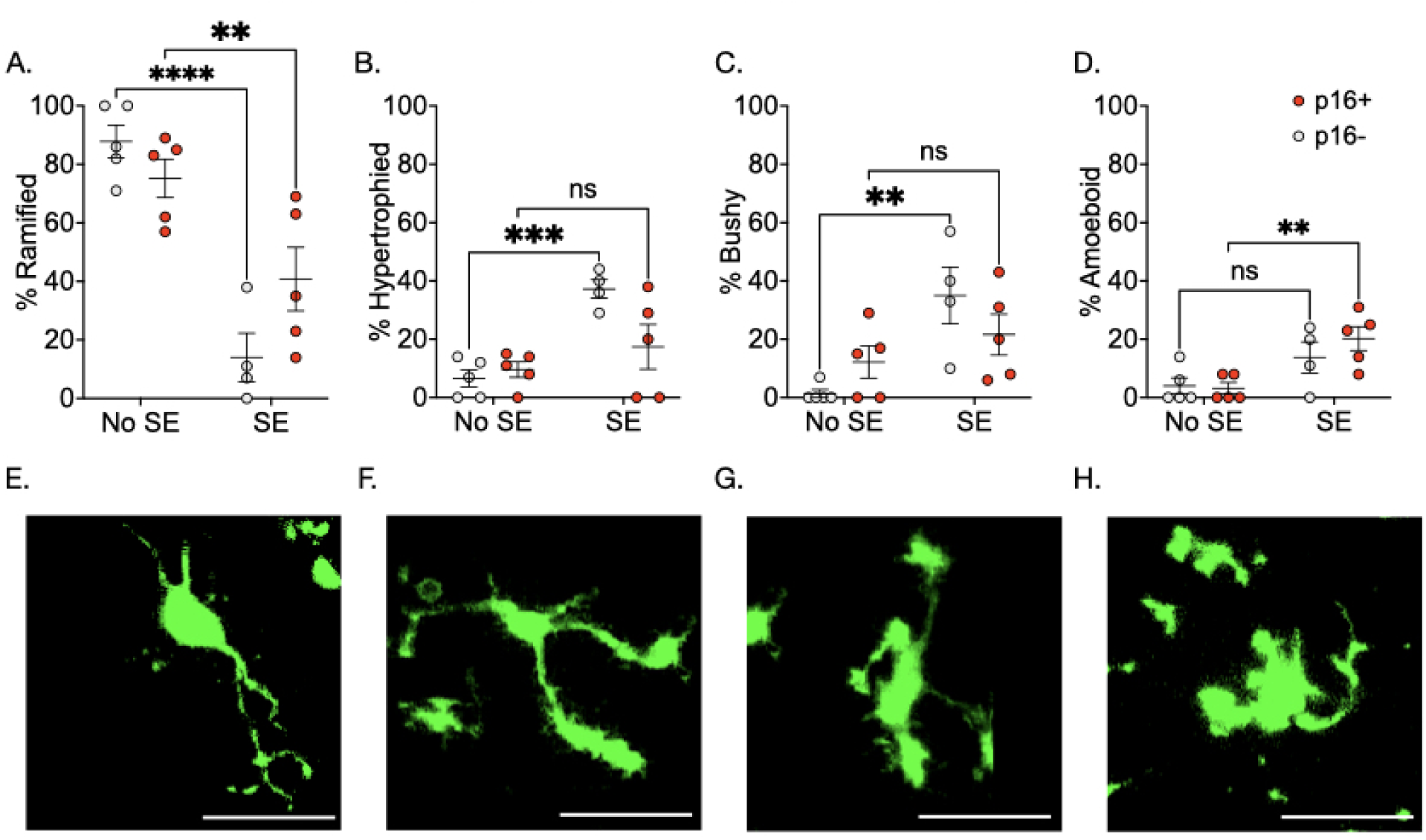
Morphology of p16 + and – microglia from *ex vivo* hippocampal slices. CX3CR1-eGFP x p16TdTom mice were crossed to examine p16 expression in red (TdTom) and microglia in green (eGFP). Live hippocampal slices from non-seizure controls (No SE) and chronic epilepsy (SE) conditions were imaged at 60x magnification to acquire Z-stack images of microglia. Microglia were randomly selected in each maximum projection image, while remaining blind to p16 expression, and characterized by morphological phenotype adapted from Wyatt-Johnson et al., 2017: ramified, hypertrophied, bushy, amoeboid, and rod. After characterization, microglia were unblinded to p16 expression (p16-: grey, p16+: red) in No SE and SE conditions. (A.) Percent of microglia expressing ramified morphology. Main effect of condition (p=0.0011, F(1,8)=9.612); no main effect on p16 expression (p=0.6944, F(1,8)=0.1660); interaction between condition and p16 expression (p=0.0147, F(1,8)=9.612). No SE vs. SE: for p16 negative microglia (p<0.0001, t=5.521, DF=16); for p16 positive microglia (p=0.4878, t=0.7102, DF=16). (B.) Percent of microglia expressing hypertrophied morphology. Main effect of condition (p=0.0333, F(1,8)=6.586), no main effect of p16 expression or interaction between condition and p16 expression. No SE vs SE for p16 negative microglia (p=0.5323, t=1.035, DF=16); for p16 positive microglia (p=0.1568, t=1.857, DF=16). (C.) Percent of microglia expressing a bushy morphology. Main effect of condition (p=0.0187, F(1,8)=0.5005), no main effect of p16 expression or interaction between condition and p16 expression. No SE vs SE for p16 negative microglia (p=0.4343, t=1.199, DF=16); for p16 positive microglia (p=0.0652, t=2.331, DF=16). (D.) Percent of microglia expressing an amoeboid morphology. Main effect of condition (p=0.0156, F(1,8)=9.36); main effect of p16 expression (p=0.0357, F(1,8)=6.358); interaction between condition and p16 expression (p=0.0178, F(1,8)=8.838). No SE vs SE for p16 negative microglia (p=0.6921, t=0.783, DF=16); for p16 positive microglia (p=0.0013, t=4.214, DF=16). (E.) Percent of microglia expressing a rod morphology. No main effect of condition, p16 expression, or interaction between condition and p16 expression. No SE vs SE for p16 negative microglia (p=0.6852, t=0.7937, DF=16); for p16 positive microglia (p=0.7.94, t=0.7556, DF=16). Representative images for each microglial morphological category: (F.) Ramified, (G.) Hypertrophied, (H.) Bushy, (I.) Amoeboid, (J). Rod. For all: Two-way ANOVA, Holms-Šídák’s multiple comparison test. Mean = +/- S.E.M.

**Supplementary Fig. 5.**
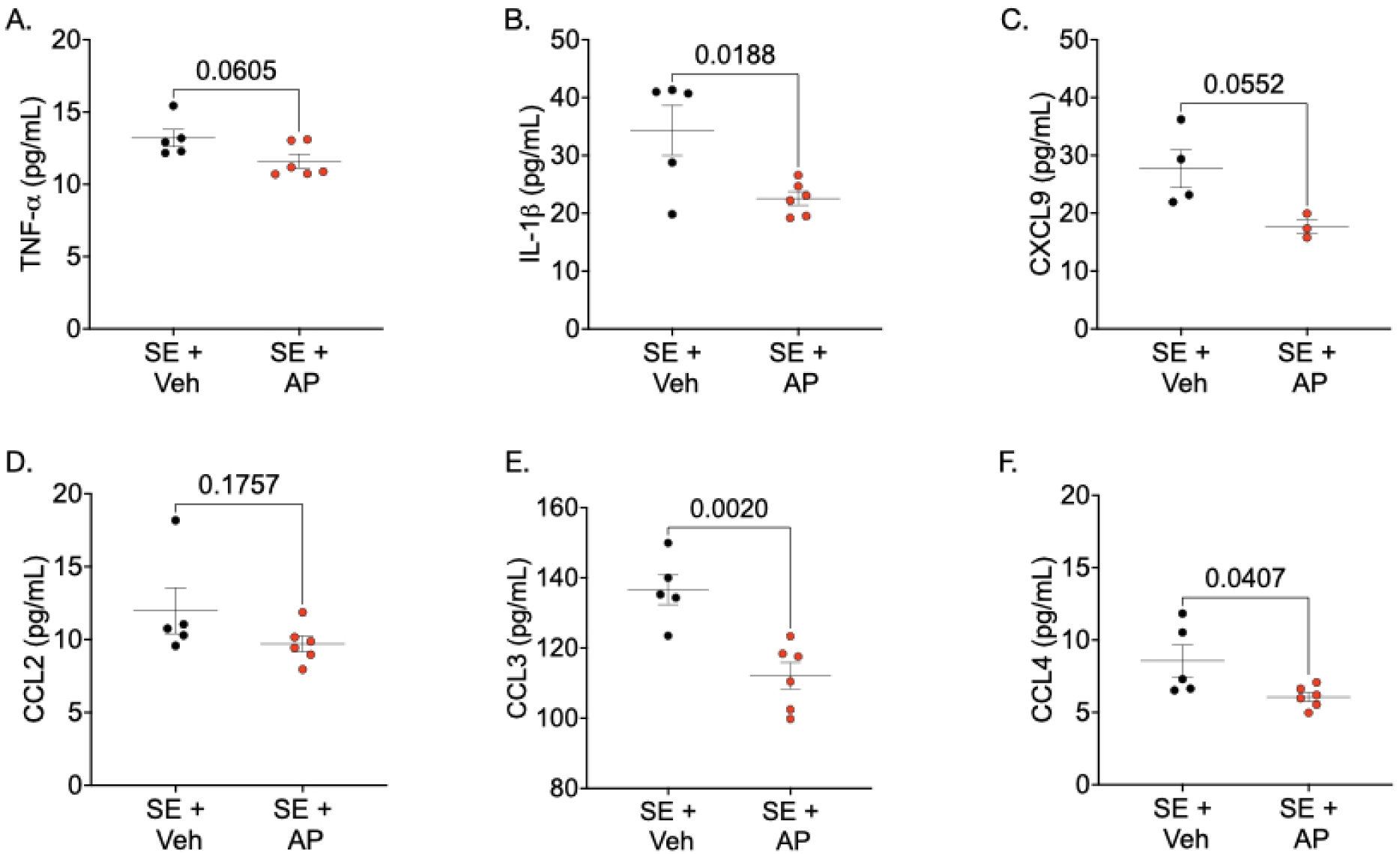
SASP expression following p16-specific senescent cell ablation in INK-ATTAC mice after epilepsy development. Protein levels of SASP factors with Legendplex multiplex mouse cytokine release panel in No SE, SE + Vehicle treated mice, SE + AP treated mice 3 months following SE. Concentration in pg/mL. (A.) TNF-α (p=0.0606, t=2.145, DF=9). (B.) IL-1β (p=0.0188, t=2.861, DF=9). (C.) CXCL9 (p=0.0552, t=2.489, DF=5). (D.) CCL2 (p=0.1757, t=1.470, DF=9). (E.) CCL3 (p=0.0020, t=4.281, DF=9) (F.) CCL4 (p=0.0407, t=2.388, DF=9). Unpaired t-test. Mean = +/- S.E.M.

**Supplementary Fig. 6.**
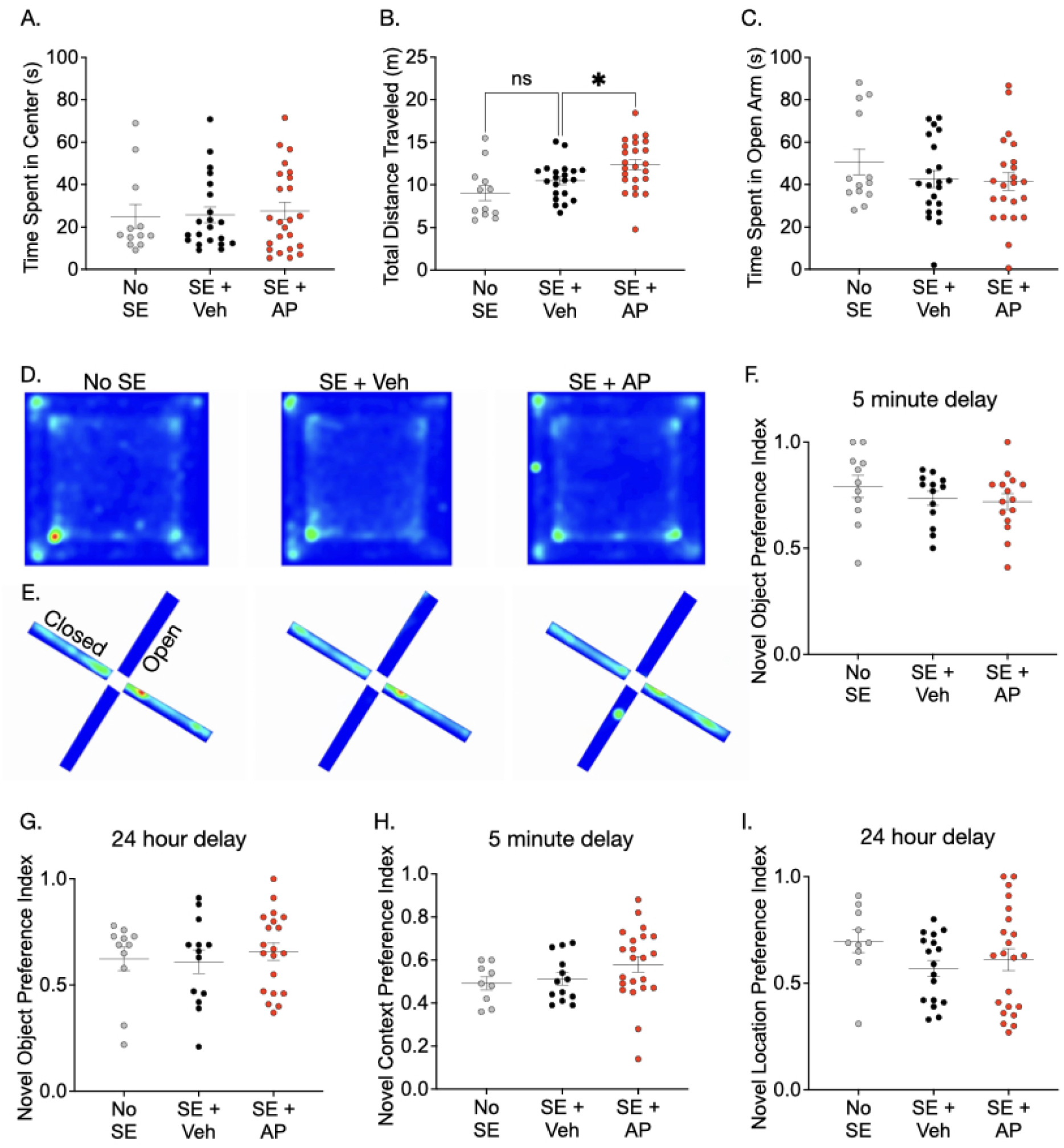
Behavioral tests following p16-specific SC ablation with AP20187 in INK-ATTAC mice. Two months after SE, after which mice developed spontaneous recurrent seizures, animals underwent a battery of behavioral tests to assess whether p16-specific SC ablation throughout epileptogenesis impacted exploratory behavior and recognition memory. (A.) No differences in time spent in the center for OF test (p=0.9094, F(2,54)=0.09515). (B.) Distance traveled during OF (p=0.9324, F(2,52)=0.0701). No SE vs. SE + Veh: p=0.1619, t=1.418, DF=54; SE + Veh vs. SE + AP: p=0.0494, t=2.306, DF=54. (C.) No differences time spent in open arm during EPM (p=0.3932, F(2,54)=0.9499). (D.) Representative heat map of animal behavior during OF: No SE (left), SE + Veh (center), SE + AP (right). (E.) Representative heat map of animal behavior during EPM: No SE (left), SE + Veh (center), SE + AP (right). (F.) No differences in NORT with a 5 minute delay before probe (p=0.4516 F(2,36)=0.8127). (G.) No differences in NORT with a 24 hour delay before probe (p=0.7563, F(2,41)=0.2812). (H.) No differences in NCRT with a 5 minute delay before probe (p=0.2122, F(2,41)=1.61). (I.) No differences in novel object location test (NOLT) with a 24 hour delay before probe (p=0.2891, F(2,46)=1.275). For all: One-way ANOVA, Holms-Šídák’s multiple comparison test. Mean = +/-S.E.M.

**Supplementary Fig. 7.**
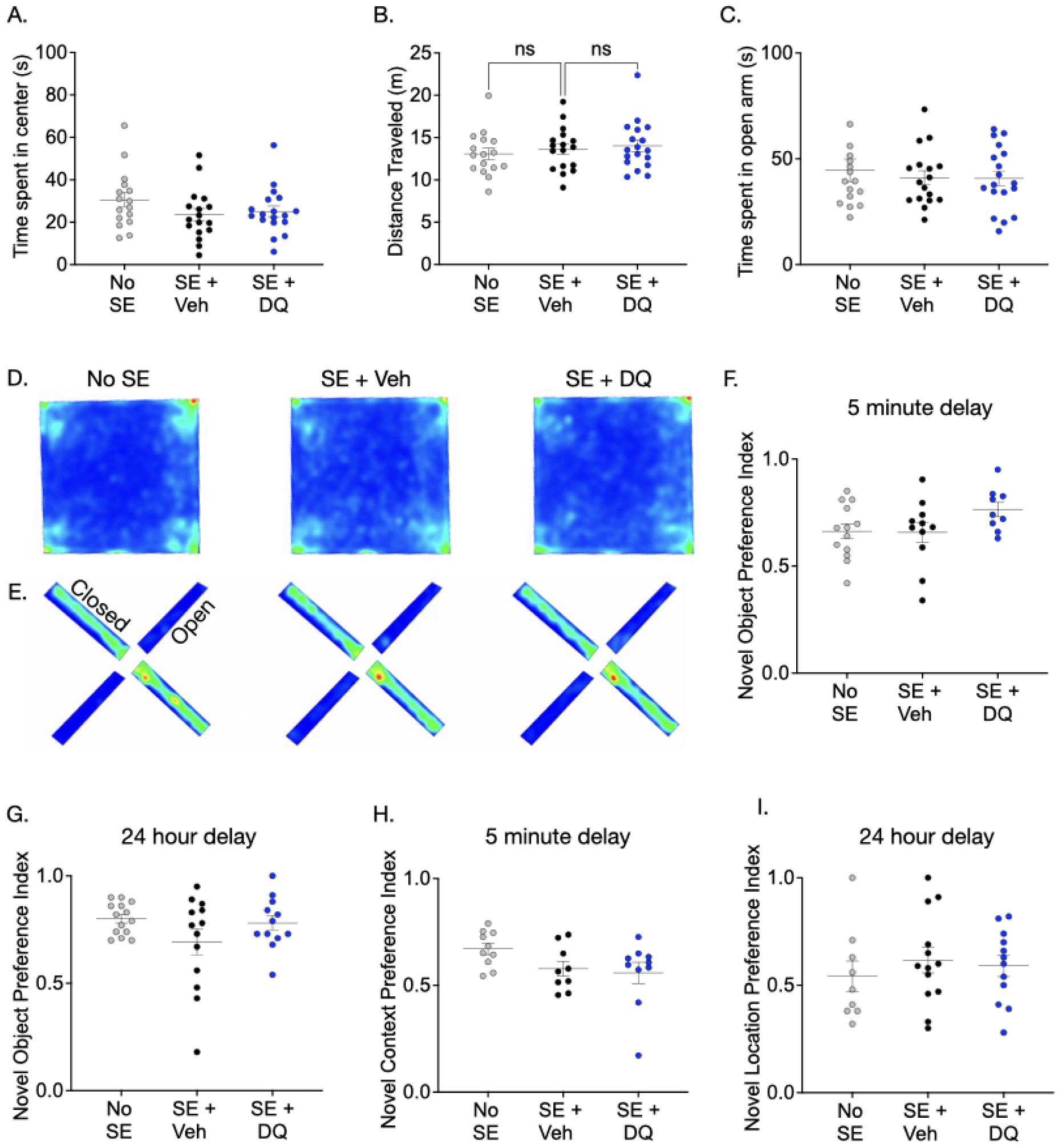
Behavioral tests following SC ablation with senolytic cocktail of DQ in C57BL/6 mice. Two months after SE, after which mice developed spontaneous recurrent seizures, animals underwent a battery of behavioral tests to assess whether DQ-mediated SC ablation throughout epileptogenesis impacted exploratory behavior and recognition memory. (A.) No differences in time spent in the center for OF test (p=0.2390, F(2,48)=1.475). (B.) Distance traveled during OF (p=0.5978, F(2,48)=0.52). (C.) No differences time spent in open arm during EPM (p=0.7690, F(2,48)=0.2641). (D.) Representative heat map of animal behavior during OF: No SE (left), SE + Veh (center), SE + DQ (right). (E.) Representative heat map of animal behavior during EPM: No SE (left), SE + Veh (center), SE + DQ (right). (F.) No differences in NORT with a 5 minute delay before probe (p=0.1497, F(2,30)=2.024) and (G.) No differences in NORT with a 24 hour delay before probe (p=0.1554, F(2, 36)=1.961). (H.) No differences in NCRT with a 5 minute delay before probe (p=0.1036, F(2, 26)=2.477). (I.) No differences in NOLT with a 24 hour delay before probe (p=0.6846, F(2,31)=0.383). For all: One-way ANOVA, Holms-Šídák’s multiple comparison test. Mean = +/- S.E.M.

**Supplementary Fig. 8.**
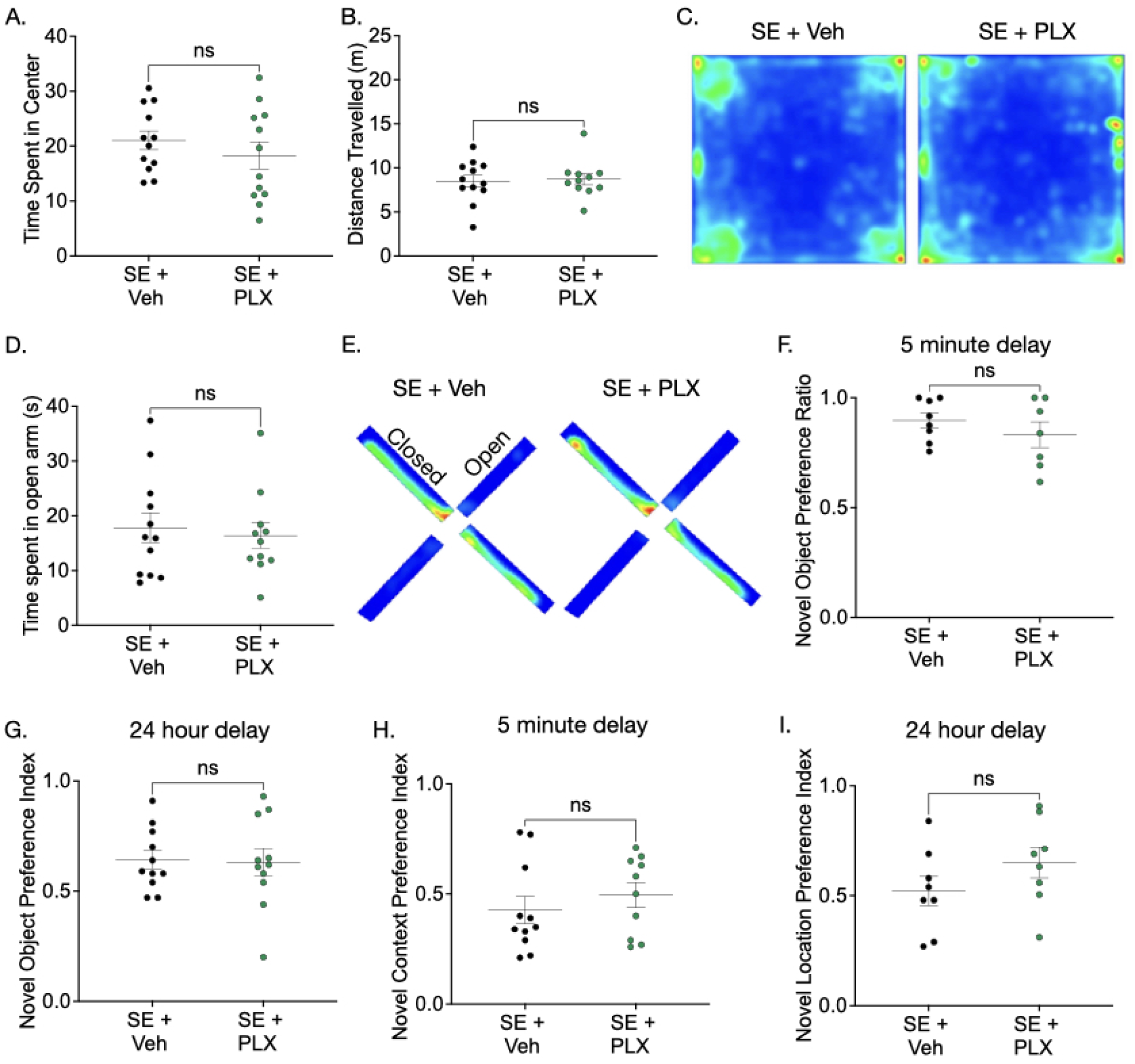
Behavioral tests following microglial ablation with PLX3397. Two months after SE, after which mice developed spontaneous recurrent seizures, animals underwent a battery of behavioral tests to assess whether microglial ablation throughout epileptogenesis impacted exploratory behavior and recognition memory. (A.) No differences in time spent in the center for Open Field (OF) test (p=0.3652, t=0.9245, DF=22). (B.) No differences in distance traveled during OF (p=0.8133, t=0.2391, DF=21) (C.) No differences in time spent in open arm during Elevated Plus Maze (EPM) (p=0.6989, t=0.3922, DF=22). (D.) Representative heat map of vehicle-treated and PLX-treated SE mice in OF. (E.) Representative heat map of vehicle-treated and PLX-treated SE mice in EPM. (F.) No differences in novel object recognition test (NORT) with a 5 minute delay before probe (p=0.332, t=1.007, DF=13). (G.) No differences in NORT probe after a 24 hour delay (p=0.8773, t=0.1564, DF=20). (H.) No differences in novel context recognition test (NCRT) with a 5 minute delay before probe (p=.4215, t=0.8216, DF=19). (I.) Novel object location test (NOLT) with a 24 hour delay before probe (p=0.2037, t=1.333, df=14). For all: Unpaired t-test. Mean = +/- S.E.M.

Supplementary Fig. 9. Representative video of microglial chemotaxis response in CA3 region from acute hippocampal slice 8 weeks post SE. Microglia extend processes towards site of 3mM ATP release. Microglia: CX3CR1-eGFP (green); p16: p16tdTomato (red).

## References

1. England MJ, Liverman CT, Schultz AM, Strawbridge LM. A Summary of the Institute of Medicine Report. Epilepsy Behav EB. 2012 Oct;25(2):266–76.

2. Ebert U, Brandt C, Löscher W. Delayed Sclerosis, Neuroprotection, and Limbic Epileptogenesis After Status Epilepticus in the Rat. Epilepsia. 2002 Jun 1;43(s5):86–95.

3. DeLorenzo RJ, Pellock JM, Towne AR, Boggs JG. Epidemiology of status epilepticus. J Clin Neurophysiol Off Publ Am Electroencephalogr Soc. 1995 Jul;12(4):316–25.

4. Maguire J. Epileptogenesis: More Than Just the Latent Period. Epilepsy Curr. 2016;16(1):31–3.

5. Kondratyev A, Gale K. Intracerebral injection of caspase-3 inhibitor prevents neuronal apoptosis after kainic acid-evoked status epilepticus. Mol Brain Res. 2000 Feb 22;75(2):216–24.

6. Didier M, Bursztajn S, Adamec E, Passani L, Nixon RA, Coyle JT, et al. DNA strand breaks induced by sustained glutamate excitotoxicity in primary neuronal cultures. J Neurosci. 1996 Apr 1;16(7):2238–50.

7. Lu JQ, Steve TA, Wheatley M, Gross DW. Immune Cell Infiltrates in Hippocampal Sclerosis: Correlation With Neuronal Loss. J Neuropathol Exp Neurol. 2017 Mar 1;76(3):206–15.

8. Parent JM, Murphy GG. Mechanisms and functional significance of aberrant seizureLinduced hippocampal neurogenesis. Epilepsia. 2008 Jun 2;49:19–25.

9. Vezzani A, Granata T. Brain Inflammation in Epilepsy: Experimental and Clinical Evidence. Epilepsia. 2005;46(11):1724–43.

10. Crowe SL, Movsesyan VA, Jorgensen TJ, Kondratyev A. Title: Rapid Phosphorylation of Histone H2A.X Following Ionotropic Glutamate Receptor Activation. Eur J Neurosci. 2006 May;23(9):2351–61.

11. Crowe SL, Tsukerman S, Gale K, Jorgensen TJ, Kondratyev AD. Phosphorylation of Histone H2A.X as an Early Marker of Neuronal Endangerment following Seizures in the Adult Rat Brain. J Neurosci. 2011 May 25;31(21):7648–56.

12. Klein P, Dingledine R, Aronica E, Bernard C, Blümcke I, Boison D, et al. Commonalities in epileptogenic processes from different acute brain insults: Do they translate? 2017;

13. Shinoda S, Schindler CK, Meller R, So NK, Araki T, Yamamoto A, et al. Bim regulation may determine hippocampal vulnerability after injurious seizures and in temporal lobe epilepsy. J Clin Invest. 2004 Apr 1;113(7):1059–68.

14. Henshall DC, Araki T, Schindler CK, Lan JQ, Tiekoter KL, Taki W, et al. Activation of Bcl-2-Associated Death Protein and Counter-Response of Akt within Cell Populations during Seizure-Induced Neuronal Death. J Neurosci. 2002 Oct 1;22(19):8458–65.

15. Castro OW, Furtado MA, Tilelli CQ, Fernandes A, Pajolla GP, Garcia-Cairasco N. Comparative neuroanatomical and temporal characterization of FluoroJade-positive neurodegeneration after status epilepticus induced by systemic and intrahippocampal pilocarpine in Wistar rats. Brain Res. 2011 Feb 16;1374:43–55.

16. De Simoni MG, Perego C, Ravizza T, Moneta D, Conti M, Marchesi F, et al. Inflammatory cytokines and related genes are induced in the rat hippocampus by limbic status epilepticus. Eur J Neurosci. 2000 Jul;12(7):2623–33.

17. Vezzani A, French J, Bartfai T, Baram TZ. The role of inflammation in epilepsy. 2011;7(1):31.

18. Vezzani A, Conti M, De Luigi A, Ravizza T, Moneta D, Marchesi F, et al. Interleukin-1β Immunoreactivity and Microglia Are Enhanced in the Rat Hippocampus by Focal Kainate Application: Functional Evidence for Enhancement of Electrographic Seizures. J Neurosci. 1999 Jun 15;19(12):5054–65.

19. Vezzani A, Pascente R, Ravizza T. Biomarkers of Epileptogenesis: The Focus on Glia and Cognitive Dysfunctions. Neurochem Res. 2017 Jul 1;42(7):2089–98.

20. Ravizza T, Gagliardi B, Noé F, Boer K, Aronica E, Vezzani A. Innate and adaptive immunity during epileptogenesis and spontaneous seizures: Evidence from experimental models and human temporal lobe epilepsy. Neurobiol Dis. 2008 Jan 1;29(1):142–60.

21. Pardoe HR, Cole JH, Blackmon K, Thesen T, Kuzniecky R. Structural brain changes in medically refractory focal epilepsy resemble premature brain aging. Epilepsy Res. 2017 Jul 1;133:28–32.

22. Hwang G, Hermann B, Nair VA, Conant LL, Dabbs K, Mathis J, et al. Brain aging in temporal lobe epilepsy: Chronological, structural, and functional. NeuroImage Clin. 2020 Jan 1;25:102183.

23. Mesraoua B, Koepp M, Schuknecht B, Deleu D, Al Hail HJ, Melikyan G, et al. Unexpected brain imaging findings in patients with seizures. Epilepsy Behav. 2020 Oct 1;111:107241.

24. Sikora E, Bielak-Zmijewska A, Dudkowska M, Krzystyniak A, Mosieniak G, Wesierska M, et al. Cellular Senescence in Brain Aging. Front Aging Neurosci [Internet]. 2021 [cited 2021 Apr 26];13. Available from: https://www.frontiersin.org/articles/10.3389/fnagi.2021.646924/full

25. Krzystyniak A, Wesierska M, Petrazzo G, Gadecka A, Dudkowska M, Bielak-Zmijewska A, et al. Combination of dasatinib and quercetin improves cognitive abilities in aged male Wistar rats, alleviates inflammation and changes hippocampal synaptic plasticity and histone H3 methylation profile. Aging. 2022 Jan 18;14(2):572–95.

26. Bussian TJ, Aziz A, Meyer CF, Swenson BL, Deursen JM van, Baker DJ. Clearance of senescent glial cells prevents tau-dependent pathology and cognitive decline. Nature. 2018 Oct;562(7728):578.

27. Zhang P, Kishimoto Y, Grammatikakis I, Gottimukkala K, Cutler RG, Zhang S, et al. Senolytic therapy alleviates Aβ-associated oligodendrocyte progenitor cell senescence and cognitive deficits in an Alzheimer’s disease model. Nat Neurosci. 2019 May;22(5):719–28.

28. Ogrodnik M, Evans SA, Fielder E, Victorelli S, Kruger P, Salmonowicz H, et al. Whole-body senescent cell clearance alleviates age-related brain inflammation and cognitive impairment in mice. Aging Cell. n/a(n/a):e13296.

29. Baker DJ, Petersen RC. Cellular senescence in brain aging and neurodegenerative diseases: evidence and perspectives. J Clin Invest. 128(4):1208–16.

30. Birch J, Gil J. Senescence and the SASP: many therapeutic avenues. Genes Dev. 2020 Dec 1;34(23–24):1565–76.

31. Crespel A, Coubes P, Rousset MC, Brana C, Rougier A, Rondouin G, et al. Inflammatory reactions in human medial temporal lobe epilepsy with hippocampal sclerosis. Brain Res. 2002 Oct 18;952(2):159–69.

32. Pernot F, Heinrich C, Barbier L, Peinnequin A, Carpentier P, Dhote F, et al. Inflammatory changes during epileptogenesis and spontaneous seizures in a mouse model of mesiotemporal lobe epilepsy. Epilepsia. 2011 Dec;52(12):2315–25.

33. Baker DJ, Wijshake T, Tchkonia T, LeBrasseur NK, Childs BG, van de Sluis B, et al. Clearance of p16Ink4a-positive senescent cells delays ageing-associated disorders. Nature. 2011 Nov 2;479(7372):232–6.

34. Baker DJ, Childs BG, Durik M, Wijers ME, Sieben CJ, Zhong J, et al. Naturally occurring p16(Ink4a)-positive cells shorten healthy lifespan. Nature. 2016 Feb 11;530(7589):184–9.

35. Ogrodnik M, Evans SA, Fielder E, Victorelli S, Kruger P, Salmonowicz H, et al. Whole-body senescent cell clearance alleviates age-related brain inflammation and cognitive impairment in mice. Aging Cell. 2021 Feb;20(2):e13296.

36. Ogrodnik M, Zhu Y, Langhi LGP, Tchkonia T, Krüger P, Fielder E, et al. Obesity-Induced Cellular Senescence Drives Anxiety and Impairs Neurogenesis. Cell Metab. 2019 May;29(5):1061–1077.e8.

37. Fatt MP, Tran LM, Vetere G, Storer MA, Simonetta JV, Miller FD, et al. Restoration of hippocampal neural precursor function by ablation of senescent cells in the aging stem cell niche. Stem Cell Rep. 2022 Feb 8;17(2):259–75.

38. Khan T, Hussain AI, Casilli TP, Frayser L, Cho M, Williams G, et al. Prophylactic senolytic treatment in aged mice reduces seizure severity and improves survival from Status Epilepticus. Aging Cell. n/a(n/a):e14239.

39. Turski W, Cavalheiro E, Bortolotto Z, Mello L, Schwarz M, Turski L. Seizures produced by pilocarpine in mice: a behavioral, electroencephalographic and morphological analysis. 1984;321(2):237–53.

40. Curia G, Longo D, Biagini G, Jones RSG, Avoli M. The pilocarpine model of temporal lobe epilepsy. J Neurosci Methods. 2008 Jul 30;172(2–4):143–57.

41. Liu JY, Souroullas GP, Diekman BO, Krishnamurthy J, Hall BM, Sorrentino JA, et al. Cells exhibiting strong p16INK4a promoter activation in vivo display features of senescence. Proc Natl Acad Sci U S A. 2019 Feb 12;116(7):2603–11.

42. Devinsky O, Vezzani A, Najjar S, Lanerolle NCD, Rogawski MA. Glia and epilepsy: excitability and inflammation. Trends Neurosci. 2013 Mar 1;36(3):174–84.

43. Altmann A, Ryten M, Di Nunzio M, Ravizza T, Tolomeo D, Reynolds RH, et al. A systems-level analysis highlights microglial activation as a modifying factor in common epilepsies. Neuropathol Appl Neurobiol. 2022 Feb;48(1):e12758.

44. Yu C, Deng X jun, Xu D. Microglia in epilepsy. Neurobiol Dis. 2023 Sep 1;185:106249.

45. Davalos D, Grutzendler J, Yang G, Kim JV, Zuo Y, Jung S, et al. ATP mediates rapid microglial response to local brain injury in vivo. 2005;8(6):752–8.

46. Avignone E, Ulmann L, Levavasseur F, Rassendren F, Audinat E. Status Epilepticus Induces a Particular Microglial Activation State Characterized by Enhanced Purinergic Signaling. J Neurosci. 2008 Sep 10;28(37):9133–44.

47. Prinz M, Jung S, Priller J. Microglia Biology: One Century of Evolving Concepts. Cell. 2019 Oct 3;179(2):292–311.

48. Kettenmann H, Hanisch UKK, Noda M, Verkhratsky A. Physiology of microglia. 2011;91(2):461–553.

49. Carvalho-Paulo D, Bento Torres Neto J, Filho CS, de Oliveira TCG, de Sousa AA, dos Reis RR, et al. Microglial Morphology Across Distantly Related Species: Phylogenetic, Environmental and Age Influences on Microglia Reactivity and Surveillance States. Front Immunol [Internet]. 2021 [cited 2021 Jul 16];12. Available from: https://www.frontiersin.org/articles/10.3389/fimmu.2021.683026/full?&utm_source=Email_to_authors_&utm_medium=Email&utm_content=T1_11.5e1_author&utm_campaign=Email_publication&field=&journalName=Frontiers_in_Immunology&id=683026

50. Jung S, Aliberti J, Graemmel P, Sunshine M, Kreutzberg GW, Sher A, et al. Analysis of Fractalkine Receptor CX3CR1 Function by Targeted Deletion and Green Fluorescent Protein Reporter Gene Insertion. 2000;20(11):4106–14.

51. York EM, LeDue JM, Bernier LP, MacVicar BA. 3DMorph Automatic Analysis of Microglial Morphology in Three Dimensions from Ex Vivo and In Vivo Imaging. eNeuro. 2018;5(6):ENEURO.0266-18.2018.

52. Pignolo RJ, Passos JF, Khosla S, Tchkonia T, Kirkland JL. Reducing Senescent Cell Burden in Aging and Disease. Trends Mol Med. 2020 Jul;26(7):630–8.

53. Xu M, Pirtskhalava T, Farr JN, Weigand BM, Palmer AK, Weivoda MM, et al. Senolytics improve physical function and increase lifespan in old age. Nat Med. 2018 Aug;24(8):1246–56.

54. Roos CM, Zhang B, Palmer AK, Ogrodnik MB, Pirtskhalava T, Thalji NM, et al. Chronic senolytic treatment alleviates established vasomotor dysfunction in aged or atherosclerotic mice. Aging Cell. 2016 Oct;15(5):973–7.

55. Xu M, Palmer AK, Ding H, Weivoda MM, Pirtskhalava T, White TA, et al. Targeting senescent cells enhances adipogenesis and metabolic function in old age. Dillin A, editor. eLife. 2015 Dec 19;4:e12997.

56. Kirkland JL, Tchkonia T. Cellular Senescence: A Translational Perspective. EBioMedicine. 2017 Jul 1;21:21–8.

57. Musi N, Valentine JM, Sickora KR, Baeuerle E, Thompson CS, Shen Q, et al. Tau protein aggregation is associated with cellular senescence in the brain. Aging Cell [Internet]. 2018 Dec [cited 2020 Dec 21];17(6). Available from: https://www.ncbi.nlm.nih.gov/pmc/articles/PMC6260915/

58. Green KN, Crapser JD, Hohsfield LA. To Kill Microglia: A Case for CSF1R Inhibitors. Trends Immunol. 2020 Sep;41(9):771–84.

59. Wu W, Li Y, Wei Y, Bosco DB, Xie M, Zhao MG, et al. Microglial depletion aggravates the severity of acute and chronic seizures in mice. Brain Behav Immun [Internet]. 2020 Jul 2 [cited 2020 Jul 6]; Available from: http://www.sciencedirect.com/science/article/pii/S0889159120302646

60. Liu M, Jiang L, Wen M, Ke Y, Tong X, Huang W, et al. Microglia depletion exacerbates acute seizures and hippocampal neuronal degeneration in mouse models of epilepsy. APSselect. 2020 Oct 1;7(10):C605–10.

61. Srivastava PK, van Eyll J, Godard P, Mazzuferi M, Delahaye-Duriez A, Van Steenwinckel J, et al. A systems-level framework for drug discovery identifies Csf1R as an anti-epileptic drug target. Nat Commun. 2018 Sep 3;9(1):3561.

62. Henning L, Antony H, Breuer A, Müller J, Seifert G, Audinat E, et al. Reactive microglia are the major source of tumor necrosis factor alpha and contribute to astrocyte dysfunction and acute seizures in experimental temporal lobe epilepsy. Glia. 2023;71(2):168–86.

63. Wyatt-Johnson SK, Sommer AL, Shim KY, Brewster AL. Suppression of Microgliosis With the Colony-Stimulating Factor 1 Receptor Inhibitor PLX3397 Does Not Attenuate Memory Defects During Epileptogenesis in the Rat. Front Neurol. 2021 Jun 3;12:651096.

64. Elmore MRP, Najafi AR, Koike MA, Dagher NN, Spangenberg EE, Rice RA, et al. Colony-Stimulating Factor 1 Receptor Signaling Is Necessary for Microglia Viability, Unmasking a Microglia Progenitor Cell in the Adult Brain. Neuron. 2014 Apr 16;82(2):380–97.

65. Engel J. Seizures and Epilepsy. OUP USA; 2013. 737 p.

66. Anovadiya AP, Sanmukhani JJ, Tripathi CB. Epilepsy: Novel therapeutic targets. J Pharmacol Pharmacother. 2012;3(2):112–7.

67. Borowicz-Reutt KK, Czuczwar SJ. Role of oxidative stress in epileptogenesis and potential implications for therapy. Pharmacol Rep. 2020 Oct 1;72(5):1218–26.

68. Martinc B, Grabnar I, Vovk T. The Role of Reactive Species in Epileptogenesis and Influence of Antiepileptic Drug Therapy on Oxidative Stress. Curr Neuropharmacol. 2012 Dec 1;10(4):328–43.

69. Citraro R, Leo A, Constanti A, Russo E, De Sarro G. mTOR pathway inhibition as a new therapeutic strategy in epilepsy and epileptogenesis. Pharmacol Res. 2016 May 1;107:333–43.

70. Engel T, Murphy BM, Schindler CK, Henshall DC. Elevated p53 and lower MDM2 expression in hippocampus from patients with intractable temporal lobe epilepsy. Epilepsy Res. 2007 Dec 1;77(2):151–6.

71. Engel T, Sanz-Rodgriguez A, Jimenez-Mateos EM, Concannon CG, Jimenez-Pacheco A, Moran C, et al. CHOP regulates the p53–MDM2 axis and is required for neuronal survival after seizures. Brain. 2013 Feb 1;136(2):577–92.

72. Burla R, La Torre M, Zanetti G, Bastianelli A, Merigliano C, Del Giudice S, et al. p53-Sensitive Epileptic Behavior and Inflammation in Ft1 Hypomorphic Mice. Front Genet [Internet]. 2018 [cited 2020 Jan 22];9. Available from: https://www.frontiersin.org/articles/10.3389/fgene.2018.00581/full

73. Flanary BE, Sammons NW, Nguyen C, Walker D, Streit WJ. Evidence that aging and amyloid promote microglial cell senescence. Rejuvenation Res. 2007 Mar;10(1):61–74.

74. Streit WJ, Xue QS. Human CNS immune senescence and neurodegeneration. Curr Opin Immunol. 2014 Aug;29:93–6.

75. Amatniek JC, Hauser WA, DelCastillo-Castaneda C, Jacobs DM, Marder K, Bell K, et al. Incidence and predictors of seizures in patients with Alzheimer’s disease. Epilepsia. 2006 May;47(5):867–72.

76. Zhang D, Chen S, Xu S, Wu J, Zhuang Y, Cao W, et al. The clinical correlation between Alzheimer’s disease and epilepsy. Front Neurol. 2022 Jul 22;13:922535.

77. Chin J, Scharfman HE. Shared cognitive and behavioral impairments in epilepsy and Alzheimer’s disease and potential underlying mechanisms. Epilepsy Behav. 2013 Mar 1;26(3):343–51.

78. Paudel YN, Angelopoulou E, Jones NC, O’Brien TJ, Kwan P, Piperi C, et al. Tau Related Pathways as a Connecting Link between Epilepsy and Alzheimer’s Disease. ACS Chem Neurosci. 2019 Oct 16;10(10):4199–212.

79. Avignone E, Lepleux M, Angibaud J, Nägerl UV. Altered morphological dynamics of activated microglia after induction of status epilepticus. J Neuroinflamm. 2015;12(1):202.

80. Sepulveda-Rodriguez A, Li P, Khan T, Ma JD, Carlone CA, Bozzelli PL, et al. Electroconvulsive Shock Enhances Responsive Motility and Purinergic Currents in Microglia in the Mouse Hippocampus. eNeuro [Internet]. 2019 Mar 1 [cited 2020 Apr 4];6(2). Available from: https://www.eneuro.org/content/6/2/ENEURO.0056-19.2019

81. Eyo UB, Peng J, Swiatkowski P, Mukherjee A, Bispo A, Wu LJ. Neuronal Hyperactivity Recruits Microglial Processes via Neuronal NMDA Receptors and Microglial P2Y12 Receptors after Status Epilepticus. 2014;34(32):10528–40.

82. Beamer E, Fischer W, Engel T. The ATP-Gated P2X7 Receptor As a Target for the Treatment of Drug-Resistant Epilepsy. 2017;11:21.

83. Beamer E, Kuchukulla M, Boison D, Engel T. ATP and adenosine-Two players in the control of seizures and epilepsy development. Prog Neurobiol. 2021 Sep;204:102105.

84. Williams-Karnesky RL, Sandau US, Lusardi TA, Lytle NK, Farrell JM, Pritchard EM, et al. Epigenetic changes induced by adenosine augmentation therapy prevent epileptogenesis. J Clin Invest. 2013 Aug;123(8):3552–63.

85. Sukigara S, Dai H, Nabatame S, Otsuki T, Hanai S, Honda R, et al. Expression of astrocyte-related receptors in cortical dysplasia with intractable epilepsy. J Neuropathol Exp Neurol. 2014 Aug;73(8):798–806.

86. Hall BM, Balan V, Gleiberman AS, Strom E, Krasnov P, Virtuoso LP, et al. Aging of mice is associated with p16(Ink4a)-and β-galactosidase-positive macrophage accumulation that can be induced in young mice by senescent cells. Aging. 2016 Jul 6;8(7):1294–311.

87. Hall BM, Balan V, Gleiberman AS, Strom E, Krasnov P, Virtuoso LP, et al. p16(Ink4a) and senescence-associated β-galactosidase can be induced in macrophages as part of a reversible response to physiological stimuli. Aging. 2017 Aug 2;9(8):1867–84.

88. Stojiljkovic MR, Schmeer C, Witte OW. Pharmacological Depletion of Microglia Leads to a Dose-Dependent Reduction in Inflammation and Senescence in the Aged Murine Brain. Neuroscience. 2022 Apr 15;488:1–9.

89. Benner B, Good L, Quiroga D, Schultz TE, Kassem M, Carson WE, et al. Pexidartinib, a Novel Small Molecule CSF-1R Inhibitor in Use for Tenosynovial Giant Cell Tumor: A Systematic Review of Pre-Clinical and Clinical Development. Drug Des Devel Ther. 2020;14:1693–704.

90. Löscher W, Stafstrom CE. Epilepsy and its neurobehavioral comorbidities: Insights gained from animal models. Epilepsia. 2023;64(1):54–91.

91. Lin YF, Wang LY, Chen CS, Li CC, Hsiao YH. Cellular senescence as a driver of cognitive decline triggered by chronic unpredictable stress. Neurobiol Stress. 2021 Nov;15:100341.

92. Neumann P, Lenz DE, Streit WJ, Bechmann I. Is microglial dystrophy a form of cellular senescence? An analysis of senescence markers in the aged human brain. Glia. 2023 Feb;71(2):377–90.

93. Zhu Y, Tchkonia T, Pirtskhalava T, Gower AC, Ding H, Giorgadze N, et al. The Achilles’ heel of senescent cells: from transcriptome to senolytic drugs. Aging Cell. 2015 Aug;14(4):644–58.

94. Jin WN, Shi K, He W, Sun JH, Van Kaer L, Shi FD, et al. Neuroblast senescence in the aged brain augments natural killer cell cytotoxicity leading to impaired neurogenesis and cognition. Nat Neurosci. 2021 Jan;24(1):61–73.

95. Matias I, Diniz LP, Damico IV, Araujo APB, Neves L da S, Vargas G, et al. Loss of lamin-B1 and defective nuclear morphology are hallmarks of astrocyte senescence in vitro and in the aging human hippocampus. Aging Cell. 2022 Jan;21(1):e13521.

96. Sano F, Shigetomi E, Shinozaki Y, Tsuzukiyama H, Saito K, Mikoshiba K, et al. Reactive astrocyte-driven epileptogenesis is induced by microglia initially activated following status epilepticus. bioRxiv. 2019 Oct 15;806398.

97. Steinhäuser C, Seifert G, Bedner P. Astrocyte dysfunction in temporal lobe epilepsy: K+ channels and gap junction coupling. Glia. 2012;60(8):1192–202.

98. Knowles JK, Xu H, Soane C, Batra A, Saucedo T, Frost E, et al. Maladaptive myelination promotes generalized epilepsy progression. Nat Neurosci. 2022;25(5):596–606.

99. Koska I, Dijk RM van, Seiffert I, Liberto VD, Möller C, Palme R, et al. Toward evidence-based severity assessment in rat models with repeated seizures: II. Chemical post–status epilepticus model. Epilepsia. 2019;60(10):2114–27.

100. Manouze H, Ghestem A, Poillerat V, Bennis M, Ba-M’hamed S, Benoliel JJ, et al. Effects of Single Cage Housing on Stress, Cognitive, and Seizure Parameters in the Rat and Mouse Pilocarpine Models of Epilepsy. eNeuro. 2019 Aug;6(4).

