## Supplemental Results for "Senescent cell clearance ameliorates temporal lobe epilepsy and associated spatial memory deficits in mice"

Human Studies. There was a significant difference in the age of patients with TLE and autopsy control patients such that the average age of TLE patients at the time of resection was nearly half of the age of patients at autopsy (33.3 +/- 13.32 vs. 65 +/- 12.80, p=0.0003, H=3.5, Mann-Whitney Test.

*Sex Differences*. All behavioral analyses were performed on sex balanced populations with a two-way ANOVA and Holms-Šídák’s multiple comparison test. Mean = +/- S.E.M.

Histology. Sex differences were not significant for p16-tdtomato, p21, or Spider B-Gal immunolabeling.

Genetic Ablation Study. No sex differences were seen for NORT (5 minute and 24 hour delay probe), NCRT, NOLT (24 hour delay probe), or BM. For NOLT with a 5 minute delay, there was no main effect of sex or treatment, however an interaction effect between sex and treatment was evident (p=0.0408, F(2,39)=3.4). There was a main effect of sex in OF (p=0.0110, F(1,43)=7.068). For EPM, a main effect of sex (p=0.0190, F(1,50)=5.875), no main effect of treatment, and an interaction effect of sex and treatment (p=0.0166, F(2,50)=4.454) were observed. There was a sex difference in EPM for SE + AP treated mice, such that males spent more time in the open arm than AP treated SE females (61.838 +/- 5.701 vs. 29.523 +/- 3.381; p=0.0005, t=4.067, DF=50). There were no sex differences in total number of seizures or average number of seizures per day. Sex differences were found in average seizure duration with a main effect of sex (p=0.0092, F(1,14)=9.104). AP-treated SE males had higher average seizure duration than females (116.49 +/- 16.07 vs. 53.04 +/- 11.98 seconds; p=0.0108, t=3.284, DF=14). There was a main effect of sex in cumulative seizure duration (p=0.0019, F(1,14)=14.63) such that males had higher cumulative seizure duration than females in both vehicle-treated (4645.92 +/- 2150.93 vs. 905.61 +/- 591.19, p=0.0481, t=2.165, DF=14) and AP-treated mice (6395.89 +/- 1424.24 vs. 792.03 +/- 547.86, p=0.0117, t=3.244, DF=14).

Senolytic Study. No sex differences were seen in OF, EPM, NORT (5 minute and 24 hour delay probe), NOLT (5 minute and 24 hour delay probe), or BM. For NCRT, SE+DQ treated males had a higher novel context preference ratio than females (0.727 +/- 0.041 vs. 0.519 +/- 0.064; p=0.0162, t=2.975, DF=33). There were no sex differences in total number of seizures, average number of seizures per day, average seizure duration, nor cumulative seizure duration.
